# Myosin VI moves on nuclear actin filaments and supports long-range chromatin rearrangements

**DOI:** 10.1101/2020.04.03.023614

**Authors:** Andreas Große-Berkenbusch, Johannes Hettich, Timo Kuhn, Natalia Fili, Alexander W. Cook, Yukti Hari-Gupta, Anja Palmer, Lisa Streit, Peter J.I. Ellis, Christopher P. Toseland, J. Christof M. Gebhardt

## Abstract

Nuclear myosin VI (MVI) enhances RNA polymerase II – dependent transcription, but the molecular mechanism is unclear. We used live cell single molecule tracking to follow individual MVI molecules inside the nucleus and observed micrometer-long motion of the motor. Besides static chromatin interactions lasting for tens of seconds, ATPase-dependent directed motion occurred with a velocity of 2 µm/s. The movement was frequently interrupted by short periods of slow restricted diffusion and increased in frequency upon stimulation of transcription. Mutagenesis and perturbation experiments demonstrated that nuclear MVI motion is independent of dimerization and occurs on nuclear actin filaments, which we also observed by two-color imaging. Using chromosome paint to quantify distances between chromosomes, we found that MVI is required for transcription-dependent long-range chromatin rearrangements. Our measurements reveal a transcription-coupled function of MVI in the nucleus, where it actively undergoes directed movement along nuclear actin filaments. Motion is potentially mediated by cooperating monomeric motors and might assist in enhancing transcription by supporting long-range chromatin rearrangements.

## Introduction

Molecular motors including the myosin family of actin-associated ATPases are enzymatic machines linked to various important cellular functions (Fili & Toseland, 2020). Besides their cytoplasmic roles, myosin motors also operate in the cell nucleus and support numerous processes. Myosin II contributes to forming the preinitiation complex and facilitates gene transcription for ICAM-1 (Li & Sarna, 2009). Nuclear myosin I and myosin V are involved in gene repositioning (Chuang et al., 2006; Dundr et al., 2007), whole chromosome translocation (Mehta, Amira, Harvey, & Bridger, 2010), transcription (A. Wang et al., 2020; Ye, Zhao, Hoffmann-Rohrer, & Grummt, 2008) and DNA damage repair (Caridi et al., 2018).

Myosin VI (MVI) is a unique member of the myosin family with the ability to move to the minus end of actin filaments (Wells et al., 1999). The motor protein contributes to transport processes including cell migration (Geisbrecht & Montell, 2002; Yoshida et al., 2004) and endocytosis (Aschenbrenner, Naccache, & Hasson, 2004; Hasson, 2003). In addition, MVI acts as load-dependent anchor in the structural maintenance of the Golgi complex (Warner et al., 2003) and in stereocilia bundles of hair cells (Avraham et al., 1995). MVI has also been found to have a role in transcription, *in vitro*, and is linked to hormone-driven gene expression (Cook, Hari-Gupta, & Toseland, 2018; Fili et al., 2020, 2017; Vreugde et al., 2006). Moreover, MVI has been found to directly interact with DNA through its Cargo Binding Domain (CBD) (Fili et al., 2017). These diverse functions are related to numerous binding partners which regulate the motor protein in monomeric and dimeric states (Fili et al., 2017). Due to the wide array of functions, it is not surprising that MVI is associated with various diseases including hypertrophic cardiomyopathy, deafness and cancer (Avraham et al., 1995; Dunn et al., 2006; Mohiddin et al., 2004; Yoshida et al., 2004).

While cytoplasmic functions of myosins are well understood and are connected to actin (Spudich & Sivaramakrishnan, 2010; Sweeney & Houdusse, 2010), little is known about the molecular mechanisms of myosin-based activity within the nucleus and the role of nuclear actin. Indeed, there is debate surrounding the nature of nuclear actin from monomers, polymers through to filaments (Plessner & Grosse, 2019). Undoubtedly, the states of nuclear actin will impact on the potential roles of myosins and there are currently no reports of actin supporting myosin-based movement in the nucleus.

To study the role of myosin VI in the nucleus, we monitored individual nuclear MVI molecules by live cell single molecule tracking and observed micrometer-long ATP-driven MVI motion on nuclear actin filaments. In addition, we found that MVI activity is required for long-range chromatin rearrangements upon transcription stimulation. Our data suggest a myosin-based cargo transportation mechanism within the nucleus, not dissimilar to that within the cytoplasm.

## Results

### Nuclear myosin VI exhibits different kinetic fractions and undergoes directed motion

We visualized MVI in live HeLa cells by an N-terminal HaloTag (Los et al., 2008) (Figure 1a and Methods). Fusion constructs were stably expressed at similar levels compared to endogenous myosin VI (Supplementary Figure 1 and Methods). As expected, TMR-HaloTag-MVI localized to the cytoplasm but also showed nuclear localization (Figure 1b and Methods)(Vreugde et al., 2006). We visualized single SiR-HaloTag-MVI molecules in the cell nucleus using HILO microscopy (Tokunaga, Imamoto, & Sakata-Sogawa, 2008) (Figure 1c and Methods). Live cell single molecule tracking (Clauß et al., 2017) of MVI revealed diverse kinetic behavior in the nucleus (Figure 1d and e and Supplementary Movie 1), suggesting MVI populates several different kinetic states. Using interlaced time-lapse microscopy (ITM)(Reisser et al., 2018), we observed that 87% of MVI molecules were fast diffusing, whereas 13% were static, probably corresponding to chromatin bound molecules (Figure 1f and Methods). In these experiments, we defined a molecule to be bound if it was confined to a radius of 160 nm for at least 100 ms. We next identified dissociation rates of bound MVI molecules from chromatin using time-lapse microscopy (Gebhardt et al., 2013) to correct for photobleaching and genuine rate identification (GRID), which allows extracting a complete spectrum of dissociation rates from single molecule fluorescence survival time distributions (Figure 1g, Supplementary Figure 2 and Methods) (Reisser et al., 2020).

**Figure 1:**
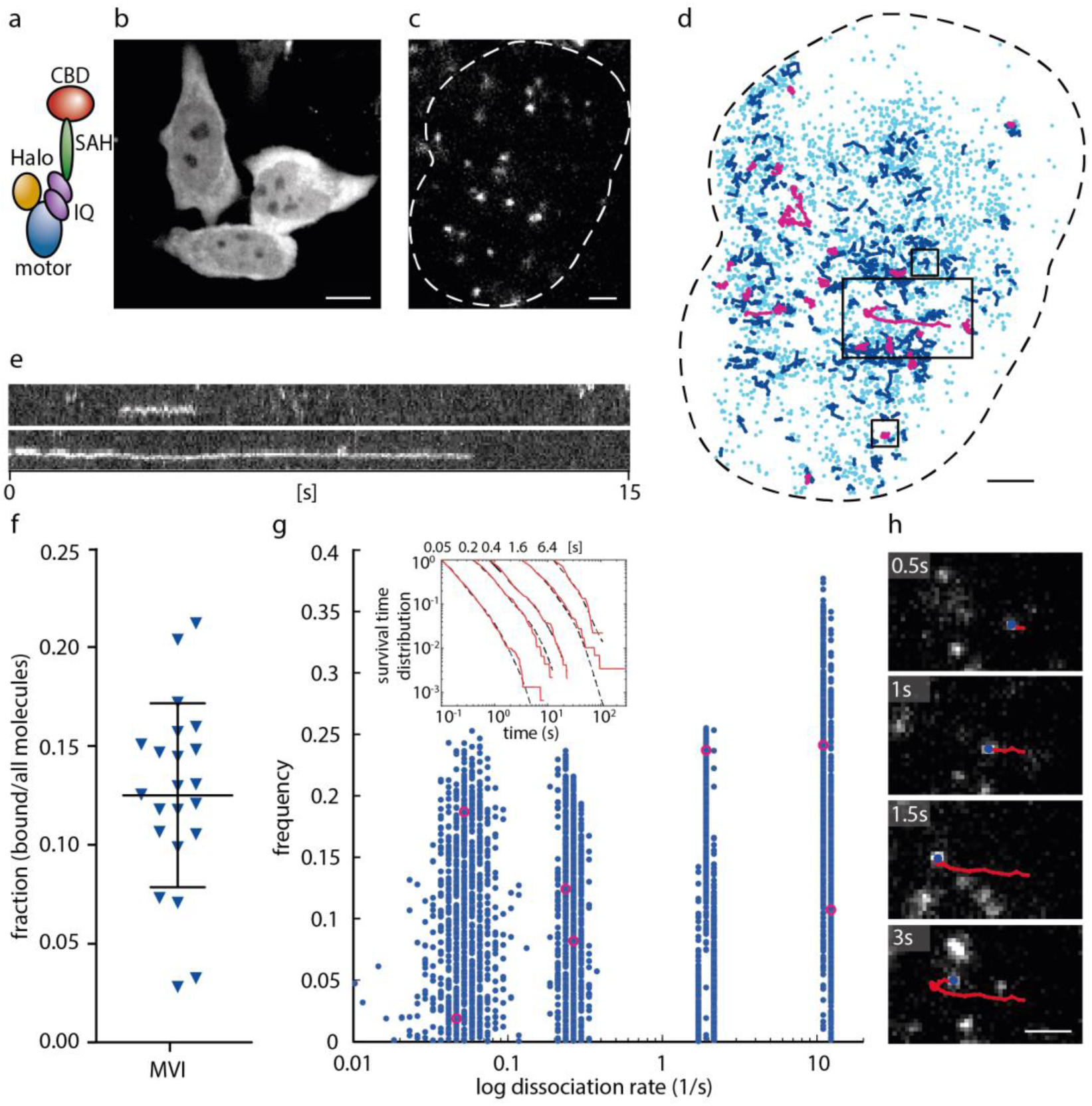
Nuclear myosin VI exhibits static chromatin binding or micrometer-long movement. (**a**) Sketch of the HaloTag-MVI construct. (**b**) Fluorescence image of a stable HaloTag-MVI HeLa cell line. (**c**) Single molecule imaging of HaloTag-MVI in the nucleus. (**d**) HaloTag-MVI detections (light blue) and tracks of HaloTag-MVI molecules lasting for > 150 ms (dark blue) or > 500 ms (magenta). (**e**) Kymographs of HaloTag-MVI tracks (upper and lower box in d). (**f**) Fraction of bound MVI molecules (mean 0.13 ± 0.05 s.d.). Data includes 30991 molecules from 22 cells. (**g**) State spectrum of HaloTag-MVI dissociation from chromatin as obtained by GRID (Reisser et al., 2020) using all data (magenta). Error estimation was obtained by resampling with 80% of the data (blue)(Methods). Inset: survival time distributions of HaloTag-MVI molecules measured at the time-lapse conditions indicated above. Dashed lines are theoretical survival time functions using the dissociation rate spectrum obtained by GRID. Data includes 6752 molecules from 32 cells. (**h**) Still images visualizing directed MVI motion (middle box in d). Scale bar is 10 µm in (b) and 2 µm in (c)(d) and (h).

We identified four distinct rate clusters, corresponding to binding time classes around 100 ms (35 ± 1.6 % spectral weight ± s.d.), 500 ms (24 ± 0.9 % spectral weight ± s.d.) 4s (19 ± 2.1 % spectral weight ± s.d.) and 19 s (21 ± 2.6 % spectral weight ± s.d.)(Methods and Supplementary Table 1). Interestingly, while the majority of bound molecules remained static at roughly the same position until dissociation, we observed a small fraction of 0.4 % of bound MVI molecules, which moved over micrometer distances (Figure 1h and Supplementary Movies 1 and 2 and Methods). Motion of MVI frequently comprised of directed runs, interrupted by pauses during which MVI apparently diffused slowly (Supplementary Movie 2). We automatically identified MVI molecules undergoing long motion by a recurrence algorithm, SPLIT (Speedy Partition of Legs In Tracks), which considered the spatial separation of molecule localizations along the track (Figure 2a, Supplementary Figure 3 and Methods). Applying this algorithm to all MVI molecules bound for at least 500 ms enabled us to distinguish statically bound MVI molecules from moving ones and to separate runs from pauses within a moving molecule track without assuming a model for pauses (Figure 2b). In a second step, we further refined the class of moving MVI molecules identified by the recurrence algorithm by including a model of slow diffusion. This enabled us to select moving MVI molecules for which the probability that a run originated from slow diffusion was below a threshold of 10^−6^ (Supplementary Figure 4 and Methods).

**Figure 2:**
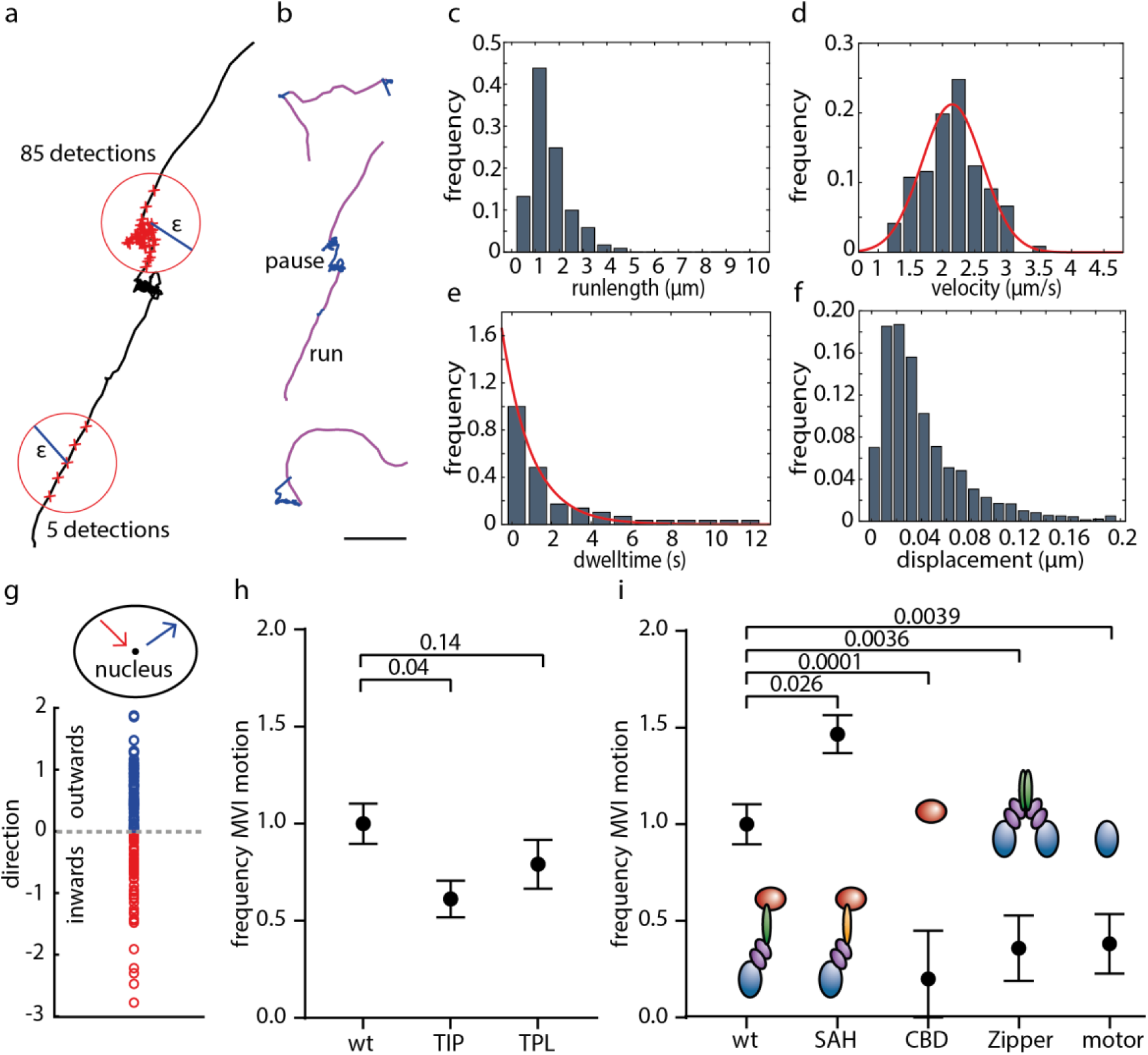
Characteristics of myosin VI motion in the nucleus. (**a**) Motion of MVI was separated into runs and pauses by a recurrence algorithm (Methods). (**b**) Examples of angled, straight or curved MVI runs (purple) with interspersed pauses (blue). Scale bar is 2 µm. Distributions of (**c**) runlength (mean 1.4 ± 0.6 µm) and (**d**) velocity (mean 2.2 ± 0.1 µm s^-1^) of MVI motion. (**e**) Distribution of dwell times (decay time 1.3 ± 0.1) and (**f**) step-size histogram (diffusion coefficient 0.01 ± 0.001 µm^2^ s^-1^) of MVI pauses. (**g**) Directionality of MVI runs (direction of end-to-end vector relative to the centre of the nucleus). (**h**) Frequency of MVI motion normalized to wild type (wt) in presence of TIP or TPL. (**i**) *Inset:* Sketch of the HaloTag-MVI truncation constructs. *Lower panel:* Frequency of MVI motion normalized to wild type (wt) of a monomeric MVI mutant (SAH), the MVI cargo binding domain (CBD), a dimeric MVI mutant (Zipper) and the MVI motor domain (motor). Frequencies in h and i were normalized to the number of detection events (n > 250,000). Detailed statistics are shown in Supplementary Table 2. Error bars denote sqrt(N), P-values are from a Chi^2^ test.

We then assessed the distribution of the kinetic parameters of the identified tracks. Runs of moving MVI molecules showed a broad distribution of run lengths, with a median run length of (1.4 ± 0.6) µm (median ± s.d.) (Figure 2c and Supplementary Figure 5). Velocities of running MVI were Gaussian distributed with a mean velocity of (2.2 ± 0.1) µm s^-1^ (mean ± s.d.) (Figure 2d), this is faster than the one measured for an artificially dimerized myosin VI construct *in vitro* (Rock et al., 2001). Pauses of moving MVI molecules had a characteristic pause time of (1.3 ± 0.1) s (decay length of exponential fit ± confidence interval) (Figure 2e) and an apparent slow diffusion coefficient of (0.01 ± 0.001) µm^2^ s^-1^ (diffusion coefficient ± sem, Methods), which is comparable to chromatin motion (Gasser, 2002)(Figure 2f). We further analyzed whether moving MVI molecules preferentially moved towards the nuclear periphery, similar to myosin I and myosin V involved in DNA repair (Caridi et al., 2018), by monitoring the direction of the end-to-end vector of a moving MVI molecule (Figure 2g). However, moving MVI did not show a preference of peripheral over central motion.

### Myosin VI long-range movement is ATPase-dependent and requires the full-length motor

To test whether motion of MVI was dependent on ATP, we compared the frequency of MVI movement in absence and presence of 2,4,6-Triiodphenol (TIP), which specifically inhibits the ATPase activity of myosin VI (Heissler et al., 2012) and was previously shown to impede transcription (Cook et al., 2018). The frequency of movement decreased to ca. 0.6 in presence of 25 µM of TIP (Figure 2h, Supplementary Figures 5 and 6, Supplementary Table 2 and Methods). Since MVI has been shown to associate with RNA polymerase II (Fili et al., 2017; Vreugde et al., 2006), we treated cells with 125 nM triptolide (TPL), which inhibits transcription by RNA polymerase II (Titov et al., 2011)(Methods). However, the frequency of MVI long-range movement events did not significantly decrease upon TPL treatment (Figure 2h and Supplementary Figure 7).

To further narrow down the molecular mechanism of MVI motion in the nucleus, we created HeLa cells stably expressing HaloTag fusions of several MVI truncations and mutants, including the isolated motor domain (motor), the CBD, an artificial dimer (Zipper) which lacks the CBD, and a monomeric version (SAH) (Methods). We compared the frequency of MVI movement between these mutants and wild type MVI (Figure 2i and Supplementary Figures 8 to 11). The motor domain and the CBD, as well as the Zipper, exhibited a clearly lower frequency of motion, while the full-length monomeric version (supplementary Movie 3) showed a 1.5-fold higher frequency. This indicates that full length MVI and, in particular, the motor and CBD-mediated interactions are required for directed movement. In contrast, dimerization is not required for motion. Additionally, we observed stepwise photobleaching during the tracks, indicating that more than one MVI contributes to this movement (Supplementary Figure 12 and Supplementary Movie 4).

### Myosin VI moves along nuclear actin filaments

Since MVI is known to move on actin filaments *in vitro* (Wells et al., 1999), and actin filaments have been observed in the cell nucleus (Plessner, Melak, Chinchilla, Baarlink, & Grosse, 2015), we tested whether manipulating actin in the nucleus would affect the frequency of MVI motion. First, we transiently expressed the actin exporter Exportin 6 (XPO6) (Stüven, Hartmann, & Görlich, 2003) in HaloTag-MVI HeLa cells (Methods), which decreased the frequency of MVI long-range movement (Figure 3a, Supplementary Figure 13 and Supplementary Movie 5). Second, we treated cells with 0.1 µM Latrunculin A (LatA) to destabilize actin filaments (Coué, Brenner, Spector, & Korn, 1987)(Methods), which reduced the frequency of MVI motion (Figure 3a and Supplementary Figure 14). Third, we transiently expressed destabilizing (R62D) or stabilizing (S14C) actin mutants (Posern, Sotiropoulos, & Treisman, 2002) fused to eYFP and a nuclear localization signal (NLS)(Methods), which reduced (R62D) or did not affect (S14C) the frequency of MVI motion (Figure 3a and Supplementary Figures 15 and 16). Overall, these results indicate a role of nuclear actin filaments in the long-range movement of MVI in the nucleus.

**Figure 3:**
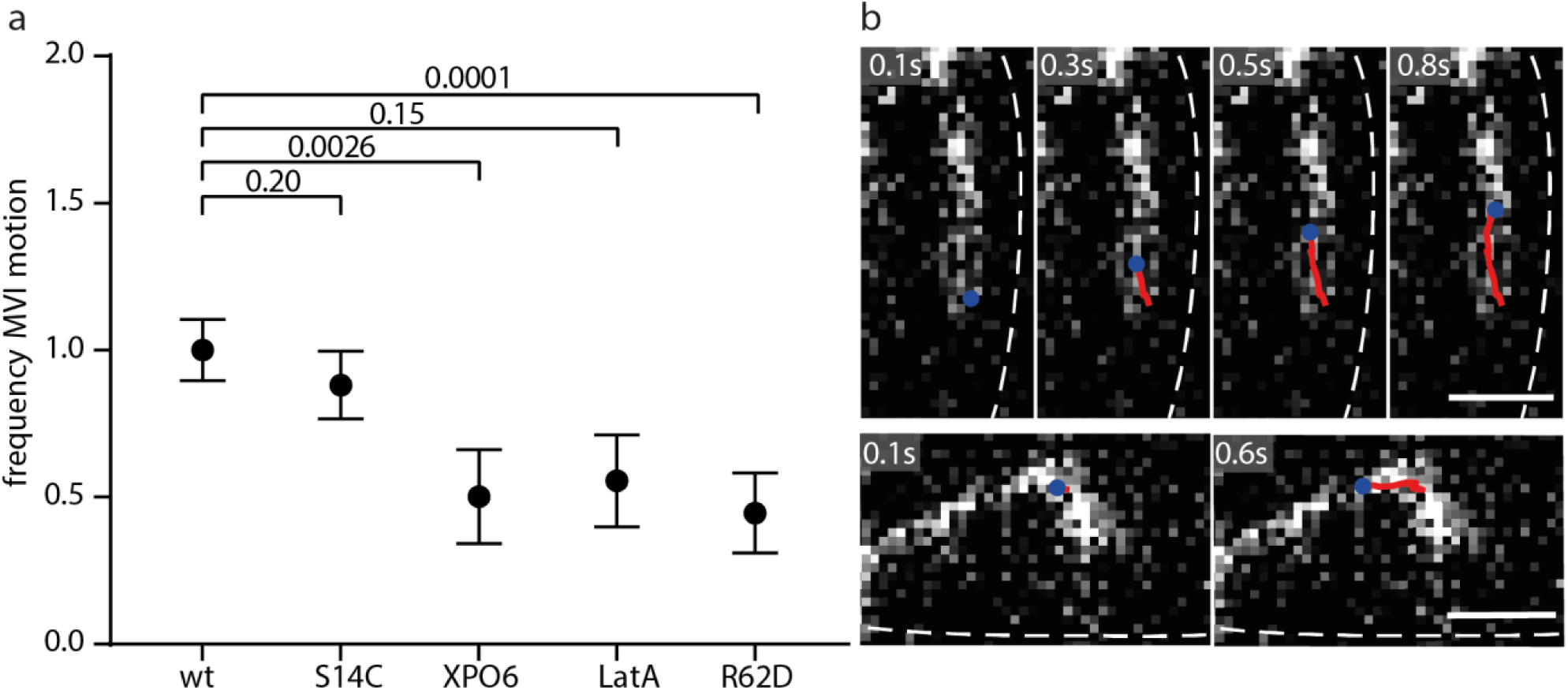
Nuclear myosin VI moves on nuclear actin filaments. (**a**) Frequency of MVI motion normalized to wild type (wt) in presence of a filament stabilizing actin mutant (S14C), an actin exporter (XPO6), Latrunculin A (LatA) or a filament destabilizing actin mutant (R62D). Frequencies were normalized to the number of detection events (n > 250,000). Detailed statistics are shown in Supplementary Table 2. Error bars denote sqrt(N), P-values are from a Chi^2^ test. (**b**) Still images visualizing co-localization of HaloTag-MVI (red) and nuclear actin chromobody (white). Scale bar is 2 µm.

To directly visualize nuclear actin filaments, we stably expressed nuclear actin chromobody (nAC)(Plessner et al., 2015) in the HaloTag-MVI HeLa cells and induced formation of transient actin filaments by 8 µM A23187 (Y. Wang et al., 2019) (Methods). Indeed, two color imaging of nAC-GFP and SiR-HaloTag-MVI revealed that MVI moved along nuclear actin filaments (Figure 3b, Supplementary Figures 17 and 18 and Supplementary Movies 6 and 7).

### Myosin VI facilitates transcription stimulated chromosome rearrangements

In order to understand what role this long-range, directed movement of MVI could play in nuclear processes, we explored hormone-stimulated transcription. It was shown previously that inhibiting MVI with TIP reduced the expression levels of estrogen receptor (ER) target genes (Fili et al., 2017). A similar effect occurred when destabilizing actin filaments with LatA (Hu et al., 2008). In addition, myosin I and actin were involved in rearranging ER target genes (Hu et al., 2008) and in chromosome movement during transcription stimulation (Chuang et al., 2006; Mehta et al., 2010). We here used the ER positive breast cancer cell line MCF7 and indeed found that MVI exhibited micrometer long movement along actin filaments in the nucleus, similar to MVI in HeLa cells (Supplementary Figures 18 and 20 and Supplementary Movie 8). Upon transcription activation of ER target genes with β-estradiol (E2) subsequent to hormone depletion with charcoal stripped FCS, the frequency of MVI motion increased (Figure 4a, Supplementary Figures 21 and 22 and Methods).

**Figure 4:**
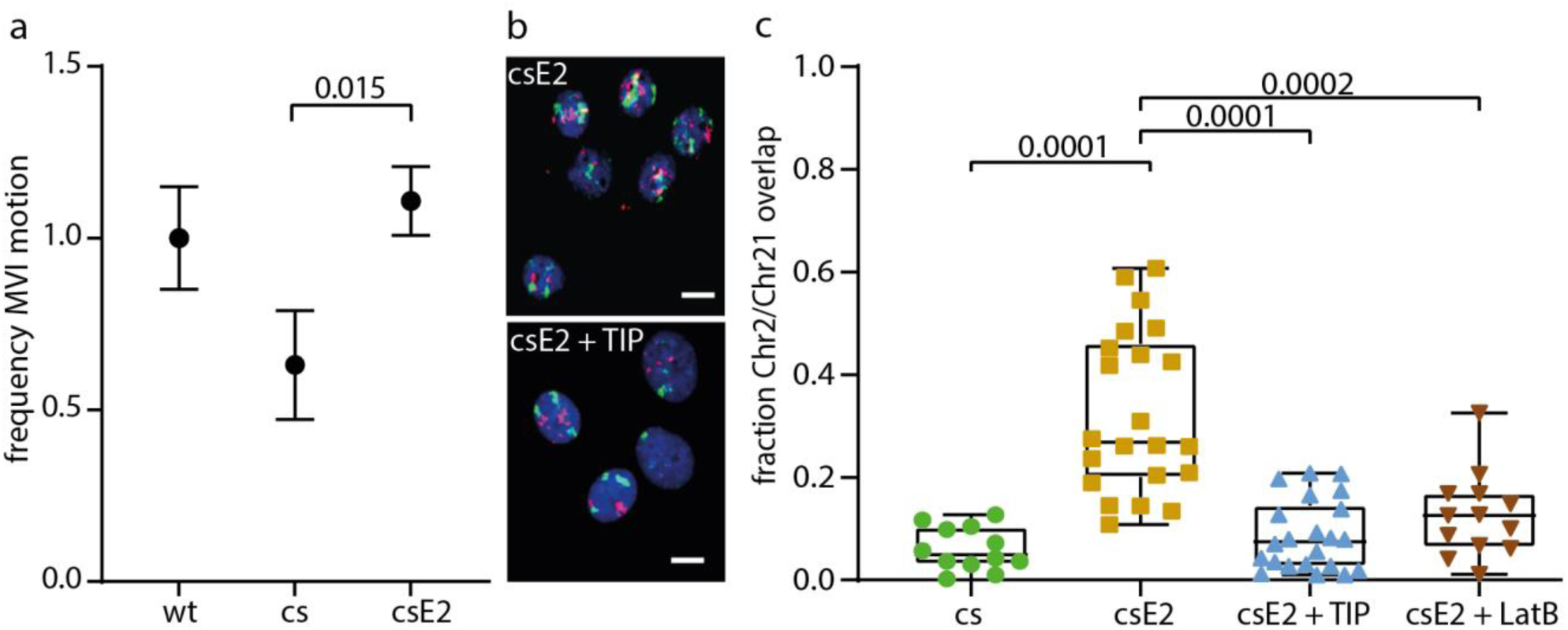
Nuclear myosin VI supports long-range chromatin rearrangements. (**a**) Frequency of MVI motion normalized to wild type (wt) in MCF7 cells in absence (cs) or presence of β-estradiol (csE2). Frequencies were normalized to the number of detection events (n > 250,000). Detailed statistics are shown in Supplementary Table 2. Error bars denote sqrt(N), P-values are from a Chi^2^ test. (**b**) Chromosome PAINT of chromosome 2 (green) and 21 (magenta) after stimulation of transcription with β-estradiol in absence (upper panel) or presence (lower panel) of the MVI inhibitor TIP in MCF7 cells. Scale bar is 10 µm. (**c**) Fraction of chromosome 2 overlapping with chromosome 21 in MCF7 cells treated 3 days with charcoal stripped FCS in absence (cs) or presence of β-estradiol (csE2), the MVI inhibitor TIP or Latrunculin B (LatB). P-values are from an unpaired two tailed t-test (n > 60 cells)

To visualise chromosome rearrangements, we used chromosome paint against chromosome 2 and 21 (Figure 4b and Methods). These chromosomes carry the genes *GREB1* and *TFF1*, respectively, and the expression of both genes is dependent upon MVI (Fili et al., 2017). After transcription activation by E2, the spatial overlap between both chromosomes increased, while treatment with TIP or Latrunculin B (LatB) inhibited chromosome rearrangements leading to spatial segregation of the chromosomes (Figure 4b and Supplementary Figure 19). This was equivalent to a transcription-inhibited state under hormone starvation conditions (Figure 4c). Overall, our experiments suggest a role of nuclear MVI and actin filaments in supporting long-range chromosome rearrangements.

## Discussion

Here, we identify a novel function for MVI in the nucleus, where it moves along nuclear actin filaments for several micrometers. Our data suggest that other nuclear motor proteins, such as RNA polymerase II, do not passively evoke this MVI motion but MVI itself actively drives the motion. First, ATPase activity of MVI is required for motion. Second, inhibition of RNA polymerase II-dependent transcription does not affect motion and third, neither the motor domain nor CBD alone allow for motion. Finally, the compromised ability of the Leucine zipper-induced MVI dimers lacking the CBD to perform this movement suggests that the movement is not solely driven by motor interactions with actin. In contrast, the monomeric MVI mutants exhibited increased mobility, indicating that movement may be driven by cooperative activity of several motors bound to cargo through the CBD, as it has been observed in vitro (Sivaramakrishnan & Spudich, 2009) and supported by our observation of more than one MVI construct on some tracks. Overall, actin-based motion of MVI in the nucleus strongly resembles its cytoplasmic function.

In total, we observed three main kinetic states that MVI can assume in the nucleus: fast diffusing, statically bound and moving. From the frequency of long-range movements, we estimate that endogenous MVI performs a run in the nucleus every 9 s when taking into account the level of overexpression of HaloTag-MVI and the labeling efficiency (Methods). Since the runs and pauses of moving MVI last on average 2 s, they will predominantly contribute to the binding time classes between 100 ms and 4 s identified by GRID. In contrast, statically bound MVI contribute to all four binding time classes including long binding events of 19 s on average. This long bound fraction of MVI might correspond to a potential tethering function of nuclear MVI, analogous to its tethering functions in the cytoplasm (Avraham et al., 1995; Warner et al., 2003). Without further study, we cannot assign the binding time classes to particular processes, binding partners or structures, although possible candidates could be the already established partners of nuclear MVI (Majewski et al., 2018), such as RNA polymerase II (Fili et al., 2017; Vreugde et al., 2006), NDP52 (Fili et al., 2017) and DNA (Fili et al., 2017).

Chromatin exhibits significant organizational order. Interphase chromosomes occupy distinct territories (M. Cremer et al., 2003; T. Cremer & Cremer, 2010; Fraser & Bickmore, 2007), are compartmentalized (Lieberman-Aiden et al., 2009) and exhibit topologically associating domains (Dixon et al., 2012) extruded by a motor function of cohesin (Davidson et al., 2019; Kim, Shi, Zhang, Finkelstein, & Yu, 2019). Moreover, long-range chromatin contacts are established, even between two different chromosomes (Beagrie et al., 2017; Lieberman-Aiden et al., 2009). Large-scale directed motion of chromatin has been visualized after transcription stimulation (Chuang et al., 2006; Dundr et al., 2007). Our data suggest that MVI supports long-range chromatin rearrangements following transcription stimulation by hormones, through a cargo transportation mechanism, as the CBD is required for motion. Based on known interactions with DNA (Fili et al., 2017), this cargo of MVI might directly be chromatin. Further supporting this notion, MVI needs to be bound to a rather static structure such as chromatin during a pause interrupting two runs, otherwise it would quickly diffuse away. Following the pause, MVI might then rebind the same or a different actin filament. In this model of long-range chromatin organization, actively moving MVI may take the role of a stir bar whisking chromatin, thereby considerably increasing the effective diffusion constant with which an otherwise spatially restricted chromosomal site is able to scan the nucleus.

Our data suggest that the role of MVI in enhancing transcription (Fili et al., 2017; Vreugde et al., 2006) is related to chromatin organization, in response to transcription stimulation. The interaction between MVI and RNA polymerase II reported previously (Fili et al., 2017; Vreugde et al., 2006) cannot be attributed to the moving activity observed here. It remains possible that MVI translocates genes to support transcription and subsequently forms interactions with RNA polymerase II at the final location of transcription. Future studies will reveal whether the statically bound fraction of MVI interacts with RNA polymerase II. Overall, we propose that nuclear MVI performs many different roles and functions, similar to its cytoplasmic pool.

## Material and Methods

### Cloning

HaloTag MVI constructs were generated as follows: MVI Motor, MVI CBD and MVI Zipper were isolated by PCR using primers containing the NLS-tag sequence (CCAAAAAAGAAGAGAAAGGTA) and then cloned into the XbaI and AscI sites of the pLV-tetO-Halo vector (Hipp et al., 2019). The monomeric MVI-SAH construct was PCR isolated similarly and cloned into the BamHI and AscI site of a modified pLV-tetO-Halo vector, which was obtained by exchanging the restriction cassette in the original pLV-TetO-Halo vector and reinserting the HaloTag N-terminal of the BamHI site. XPO6 was obtained from HeLa cell cDNA and cloned into XhoI and EcoRI sites of the pIRES-2-eGFP backbone (Doerner, Febvay, & Clapham, 2012). Supplementary Table 3 lists oligos used for cloning.

### Cell culture and creation of cell lines

All experiments were performed in HeLa cells if not indicated otherwise. HeLa cells were cultured in DMEM supplemented with 10% FBS, 1% Glutamax, 1% nonessential amino acids, and 1% sodium pyruvate. MCF7 cells were cultured in MEM supplemented with 10% FBS, 1% Glutamax and 1% nonessential amino acids. Stable cell lines were created by using a standard lentiviral production protocol (Addgene). Briefly, 293T packaging cells were transfected with psPAX2 (Addgene 12260), pMD2.G (Addgene 12259) and the transfer plasmid containing the tagged construct. Cells were stained with 2.5 µM HaloTag-TMR ligand following the manufacturer protocol (Promega) and transfected cells were sorted via their fluorescence signal (BD FACSAria III, BD, USA).

### Preparation of cells for FACS and imaging

For fluorescence activated cell sorting (FACS), cells were stained with different concentrations of TMR HaloTag ligand according to the HaloTag labeling protocol (Promega) and 50,000 cells were analyzed by flow cytometry (BDLSR Fortessa, BD, USA).

For confocal imaging, cells were stained with 2.5 µM HaloTag-TMR ligand following the manufacturer protocol (Promega) and imaged in OptiMEM at a spinning disc microscope.

For single molecule imaging, cells were seeded on Delta-T glass bottom dishes (Bioptechs, Pennslyvania, USA) and stained with SiR-HaloTag ligand (Lukinavičius et al., 2013)(kindly provided by Kai Johnson, MPI, Heidelberg, Germany) prior to imaging with a final concentration of 31 pM according to the HaloTag protocol (Promega). For binding time measurements, cells were stained with PA-JF-647 (kindly provided by Luke Lavis, Janelia Research Campus, Ashburn, USA) with a final concentration of 100 nM according to (Grimm et al., 2016). Cells were imaged in OptiMEM.

#### Additional cell treatments

For measurements in presence of TIP (Sigma), cells were treated with 25 µM 4h prior to the measurements. For measurements in presence of TPL, cells were treated with 125 nM TPL (Sigma) for 1h prior to the measurements. For measurements in presence of overexpressed XPO6, cells transiently expressing XPO6 were selected by the fluorescence of eGFP, which was co-expressed with the IRES2 system. For measurements in presence of transiently expressed eYFP-actin constructs, expressing cells were selected by the fluorescence of eYFP. For measurements in presence of LatA (Sigma), cells were treated with 100 nM LatA 5 min prior to the measurments and measurements were performed up to 20 min after treatment with LatA. For starvation and hormone stimulation experiments in MCF7 cells, cells were starved in charcoaled stripped medium for 48 h and stimulated for 5 minutes with 100 nM β-estradiol (Sigma) prior to the measurements. Stimulated cells were measured for up to 30 min after stimulation. Nuclear actin filaments were visualized with a GFP tagged nuclear actin chromobody (Chromotek) and filaments were induced with 8 µM Ionphore A23187 (Sigma) (Y. Wang et al., 2019) prior to the measurements.

### Western Blot

We loaded 25/15 µg (HeLa/MCF7) of total protein isolated from a confluent 10 cm culture dish. We used a standard Western Blot protocol with antibodies against MyoVI (M0691, Sigma, Germany, dilution 1:500) and γ-tubulin (ab11316, abcam, UK, dilution 1:200,000). Bands were visualized with a secondary antibody (Invitrogen A24527, USA) via alkaline phosphatase and quantified with ImageLab 5.0 (ChemDoc MP, BioRad, USA). The integrated intensity was used to estimate the expression level of tagged MVI compared to the endogenous level.

### Single molecule microscopy

The single molecule fluorescence microscope was home built as described previously (Reisser et al., 2018). In brief, a 405 nm laser (Laser MLD, 299 mW, Solna, Sweden), a 488 nm laser (IBEAM-SMART-488-S-HP, 200 mW, Toptica, Gräfelfing, Germany) and a 638 nm laser (IBEAM-SMART-640-S, 150 mW, Toptica, Gräfelfing, Germany) were collimated and used to set up a highly inclined illumination pattern on a conventional fluorescence microscope (TiE, Nikon, Tokyo, Japan) using a high-NA objective (100×, NA 1.45, Nikon, Tokyo, Japan). Emission light had to pass a multiband emission filter (F72–866, AHF, Tübingen, Germany) and was subsequently detected by an EMCCD camera (iXon Ultra DU 897U, Andor, Belfast, UK) with 50 ms integration time, equaling the illumination time. For time-lapse imaging, dark-times were controlled by an AOTF (AOTFnC-400.650-TN, AA Optoelectronics, Orsay, France). Temperature control was realized by the Delta-T system (Bioptechs, Pennsylvania, USA).

### Single molecule imaging and analysis

Cells were prepared as described above and kept in OptiMEM during imaging for up to 90 minutes per dish.

#### Time lapse illumination

Time lapse illumination and detection of single molecules was done as described earlier (Clauß et al., 2017). In brief, consecutive frames were recorded with different dark times to impede premature photo bleaching. The dark time between two consecutive frames was varied between 0.05 s and up to 6.4s for individual time lapse conditions (Figure 1 g). Single molecules were detected by their fluorescence intensity compared to background fluorescence and localization was performed by a 2D Gaussian fit. HaloTag-MVI molecules were considered bound if they did not leave a radius of 240 nm for 3 (50 ms and 200 ms time lapse condition) and 2 (other time lapse conditions) consecutive frames. The resulting fluorescence survival times were analysed with GRID.

#### GRID

We analyzed fluorescence survival time histograms resulting from time lapse imaging using GRID (Reisser et al., 2020). GRID is an approach for the genuine identification of decay processes contained in survival histograms. In brief, it fits a superposition of 0<N<500 exponential functions with fixed decay rate constants to the binding time histogram and returns the amplitudes corresponding to each of the fixed decay rates. An error estimation is obtained by 500 resampling runs using each run a random subset of 80% of the data (Efron, 1979). Binding times were calculated from the inverse of the dissociation rates. In case of clusters of dissociations rates, we calculated the mean dissociation rate before taking the inverse. The spectral weight is the sum over all amplitudes in one cluster obtained by resampling. The error of the spectral weight is the standard deviation of the resampling results.

#### ITM

To assess the fraction of bound molecules, we used the illumination scheme of interlaced time lapse microscopy (ITM) (Reisser et al., 2018). In an ITM measurement two camera frames are recorded followed by a dark time. After the dark time this pattern is repeated. We performed ITM measurements with an illumination time of 10 ms. The dark time was chosen such that short bound molecules vanished, while long bound molecules were still bound after the dark time. Accordingly, molecules that appeared in images before and after the dark time were categorized as long bound while molecules that only appear before or after the dark time were categorized as short bound. Molecules that were only detected in one frame were classified as diffusive.

#### Calculations and Estimations

The diffusion coefficient (D) was calculated from the mean displacement <r^2^> as 2D isotropic diffusion, with t reflecting the integration time.

Recurrence analysis showed that only a small fraction of all molecules (total-#events) is involved in MVI motion (motion-#events). We calculated this fraction by dividing total-#events by motion-#events (Supplementary table 2). Recurrence analysis of 179 movies (15 s/movie, total time = 2685 s) from 81 cells, identified 92 MVI motion events (#motion, Supplementary table 2). Considering that we have a HaloTag-MVI overexpression of 1.25 (Supplementary Figure 1a), and a HaloTag labelling efficiency of 1/4 (Supplementary Figure 1b), we estimated the frequency of endogenous MVI motion by multiplying the detected MVI motion frequency (#motion/total time) with a factor of 3.2 (1/(labelling efficiency*overexpression)), revealing a motion frequency of 0.11 #motion/s, which equals an occurrence rate of 1 event every 9s.

### Heuristic classification of directed motion (SPLIT)

#### Preprocessing before recurrence analysis of tracks

Due to localization errors, we applied a sliding average method to the tracks. We averaged x- and y-coordinates separately, using the left and right neighboring points.

#### Classification

We developed an algorithm to distinguish between different modes of motion. We first decide for each point of a track if it is in a directed or diffusive state of motion. Next, we use a data compression method to eliminate false classifications. As a last step we filtered out tracks that did not allow a confident separation between different states of motion.

Since we have no a priori information about what kind of motion to expect, our algorithm heuristically separates different modes using two characteristics: (i) Number of points in close proximity and (ii) angle between two jumps.

Let us consider a track *Tr* consisting of *N* coordinate vectors. The *n* -th vector of the track contains the coordinates 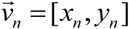. To evaluate characteristic (i) we used the recurrence-plot (Supplementary Figure 3). The recurrence plot is constructed according to the rule

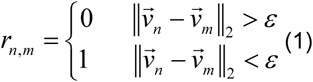

This rule implies that the recurrence plot of a track with *N* coordinate vectors is a *N* × *N* matrix, which is symmetric with respect to the diagonal. When multiple coordinates are in close proximity and separated by less than the threshold *ε* from each other, the recurrence matrix contains clusters. We quantified these clusters by summing over the diagonal in the recurrence plot to obtain the projection *h*

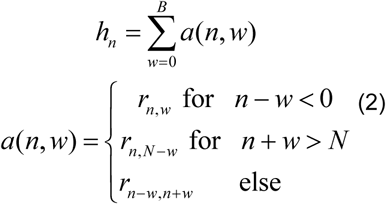

where we sum over *B* elements on the diagonal. For clarity the path of summation-indices is depicted in Supplementary Figure 3.

For the second characteristic (ii), we calculated the vectors

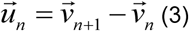

We call the angles between the vectors 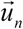 and 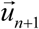, *ϑ*_*n*_. We classified the motion as directed if *ϑ*_*n*_ < *ϑ*_*direct*_ and as non-direct if the angle was larger than *ϑ*_*n*_ > *ϑ*_*non*−*direct*_. We observed that the angle is strongly affected by localization errors. To obtain a more robust quantity we took the average angle of three subsequent points in the track.

We unify criteria (i) and (ii) in a classification into directed and non-directed movement according to the rules

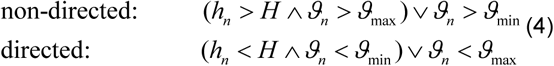

where we introduce the threshold *H* to distinguish between directed and non-directed motion. According to the above equation, the recurrence criterion (i) makes a soft decision that can be revoked be the angle criterion (ii).

#### Compression and averaging of classification results

The above procedure yields a sequence of classifications into directed and non-directed movement. We compressed this sequence into tupels. A tuple consisted of a movement type and the number of adjacent time points the molecule adopted this type of movement. To filter out outliers in classification we deleted tuples with a duration smaller than *τ* _min_ and added their value to the largest neighboring tuples.

#### Filtering of tracks

Since diffusion is a stochastic process, diffusing tracks may contain straight pieces of movement with long jump distances. Such parts of a track would be falsely classified as directed motion. To exclude these statistical outliers, we calculated the probability that a segment of apparent directed movement was caused by diffusion. If this probability was higher than 1e-6 for one sequence of directed motion, we excluded the track from further analysis.

#### Choice of parameters

Above we have introduced several parameters such as the summation border *B*, the threshold *H*, the recurrence plot parameter *ε* and the minimum and maximum angles *ϑ*_*non*−*direct*_ and *ϑ*_*direct*_. We now give a reasoning of how we chose the values of these parameters. Directed movement in a circle-shape will generate many points in close proximity to each other. To prevent that such tracks were wrongly classified as non-directed we chose a summation border of five. Once a coordinate vector in the track had more or equal than three neighbors in close proximity we classified this point as non-directed. We chose *ε* = 0.4μm. Assuming a diffusion coefficient of chromatin bound molecules of 0.01μm^2^s^-1^ (Akhtar & Gasser, 2007) the chance that the molecule escapes that radius goes to zero.

The angles make a hard decision on whether the motion is directed. We therefore chose values that clearly indicate directed or non-directed motion: *ϑ*_*non−direct*_ = 45° and *ϑ*_*direct*_ = 5°

### 2D Fish Analysis

MCF7 cells were cultured and cells were starved and then stimulated with E2, as described above. They were additionally treated with 25 µM TIP for 4 hrs and 100 nM Latrunculin B for up to 1 hr. Once all treatments had been completed cells were washed with PBS and then harvested. The cells were then centrifuged at 500 x g for 10 mins at RT. The cells were resuspended in 0.075 M KCl and incubated at 37°C for 20 mins before repeating the centrifugation. The cells were then resuspended in Fix solution (3 parts methanol: 1 part glacial acetic acid) and centrifuged at 500 x g for 10 mins. This process was repeated three times to wash the cells into the Fix solution. A Vysis HYBrite Hybridization System was preheated to 37°C. 10µL of cells in Fix solution were dropped directly onto a microscope slide followed by 10µl of Fix solution only. The slide was left to dry at RT and then immersed into 2x saline-sodium citrate solution (SSC) for 2 mins at RT without agitation. The slides were then dehydrated in an ethanol series (2 mins in 70%, 85% and 100% ethanol) at RT. They were then air dried and placed on the hybridisation system at 37°C. Cover slips were prepared with 10µL of the chromosome paint (Chromosome 2 and 21 Cytocell) and placed onto the slides. The coverslips were sealed using rubber cement and placed back onto the hybridisation system at 75°C for 2 mins. The samples were left within a hybridisation chamber overnight in the dark at 37°C. The rubber cement and coverslips were removed and the slides were immersed into 0.4xSSC pH 7.0 at 72°C for 2 mins, without agitation. The slides were drained and then placed into 2xSSC supplemented with 0.05% Tween-20 at RT for 30 secs. The slides were air-dried and 10µl of DAPI was placed onto each sample and a coverslip was placed and sealed onto the samples. Images were taken using an Olympus BX53 with a 40x oil-immersion objective. Over 60 nuclei per condition were then analysed using the nuclear morphology analysis software, (https://bitbucket.org/bmskinner/nuclear_morphology/wiki/Home)(Skinner et al., 2019).

## Code availability

The code for the recurrence algorithm to analyse tracks of myosin VI is freely available. A MatLab version of the code is available at https://gitlab.com/GebhardtLab/SPLIT.

## Data availability

Data supporting the findings of this manuscript will be available from the corresponding author upon reasonable request. All raw single particle tracking data will be freely available in Matlab and csv file format after publication.

## Acknowledgements

We thank Robert Grosse and Matthias Plessner for comprehensive help with nuclear actin filament visualization. We thank Astrid Bellan-Koch for help with Western Blots. SiR dye was kindly provided by Kai Johnsson (Max Planck Institute for Medical Research, Heidelberg, Germany). PA-JF646 was kindly provided by Luke Lavis (Janelia Research Campus, Ashburn, USA). pIRES-2-eGFP was kindly provided by David E. Clapham (Boston Children’s Hospital, Boston, MA, USA). The work was funded by the European Research Council (ERC) under the European Union’s Horizon 2020 Research and Innovation Program (No. 637987 ChromArch to J.C.M.G.), the German Research Foundation (SPP 2202 GE 2631/2-1 and GE 2631/3-1 to J.C.M.G.) and UKRI-Medical Research Council (MR/M020606/1 to C.P.T.). J.C.M.G. acknowledges support by the Collaborative Research Center 1279. The authors thank the Ulm University Center for Translational Imaging MoMAN for its support.

## Author contributions

C.P.T. and J.C.M.G. conceived the study. A.G-B., A.W.C., C.P.T. and J.C.M.G. designed experiments. C.P.T, N.F., A.P. and L.S. designed and cloned myosin VI mutants. A.G-B. created cell lines. A.G-B. performed and analyzed single molecule tracking experiments. J.H. developed the recurrence algorithm. J.H., T.K. and J.C.M.G. contributed to data analysis. A.W.C, Y.H-G. and P.J.I.E. performed chromosome paint experiments. A.W.C. and C.P.T. analyzed chromosome paint experiments. C.P.T. and J.C.M.G. supervised the study. J.C.M.G., A.G-B. and C.P.T. wrote the manuscript with comments from all authors.

