## Supplementary Information for "Myosin VI moves on nuclear actin filaments and supports long-range chromatin rearrangements"

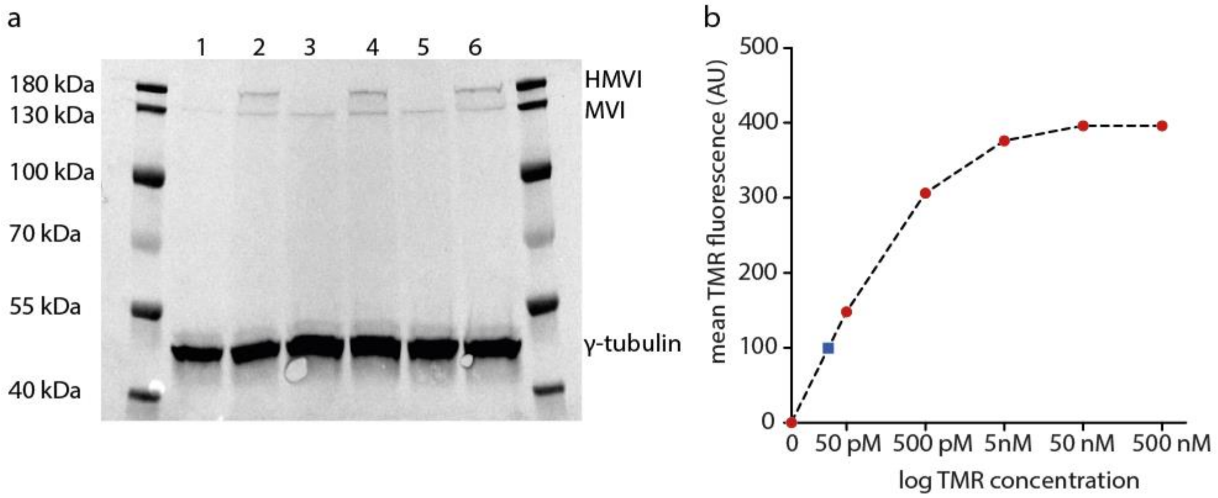


**Supplementary Figure 1:** Quantification of overexpression and labeled fraction of HaloTag-myosin VI. (**a**) Western Blot of endogenous myosin VI (MVI) and HaloTag-myosin VI (HMVI) expression in HeLa cells (lane: 1, 3 and 5), and a cell line stably expressing HaloTag-myosin VI (lane: 2, 4 and 6) (antibody M0691 Sigma, concentration 2 µg/ml). γ-tublin was used as loading control (antibody ab11316 abcam, concentration 5 ng/ml). Lanes 1/2, 3/4 and 5/6 are biological replicates. HaloTag-myosin VI was overexpressed by 1.25 ± 0.11 compared to endogenous myosin VI. (**b**) Titration of TMR-HaloTag ligand. Cells expressing HaloTag-myosin VI were labeled with TMR-HaloTag ligand at various concentrations (red dots) (Methods). The mean TMR fluorescence intensity of 50,000 cells at every concentration was determined by flow cytometry. Single molecule tracking experiments where performed with a SiR-HaloTag ligand at a concentration of 31 pM (blue square). Assuming similar labeling efficiencies of TMR-HaloTag ligand and SiR-HaloTag ligand, ca. 1/4 of ectopically expressed HaloTag-MVI molecules were labeled with SiR-HaloTag ligand.

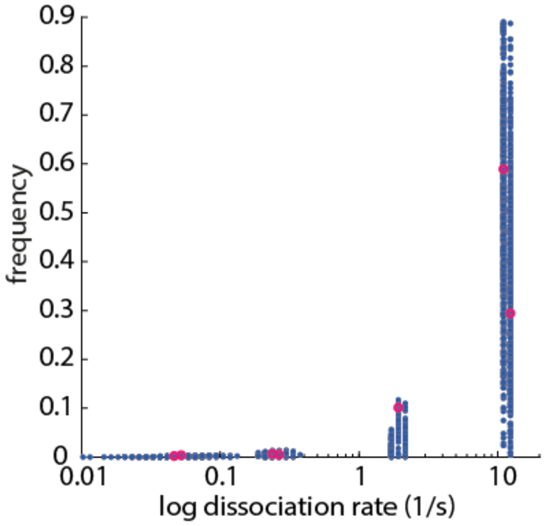


**Supplementary Figure 2:** Event spectrum of HaloTag-MVI dissociation from chromatin as obtained by GRID using all date (magenta). Error estimation obtained by resampling of 80% of the data (blue) (Supplementary Table 1 and Methods).

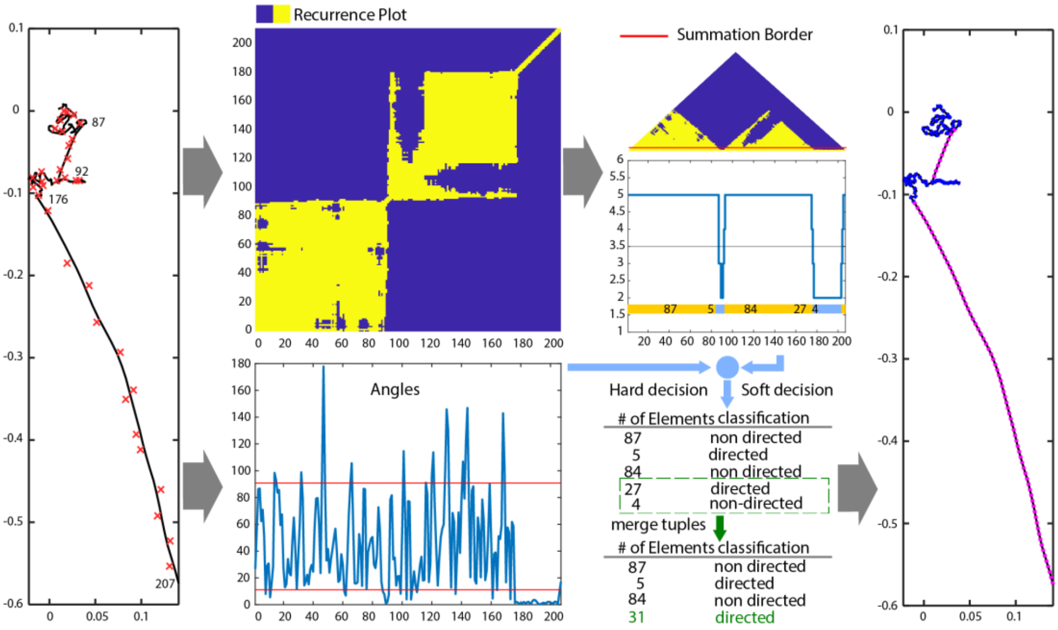


**Supplementary figure 3:** Analysis of a simulated track. We first applied a sliding average on the track coordinates to smoothen detection inaccuracies (red crosses: detections, black line: smoothened track). We then calculated the recurrence plot according to equation (1) and performed a soft decision for directed and non-directed movement following equation (2). We combined this classification with the angle criterion that can override the recurrence classification if it clearly indicates directed or non-directed movement (equation (4)). The resulting sequence of classifications in the track was compressed to tuples. To eliminate outliers, short tuples were merged with larger ones. See text for further details.

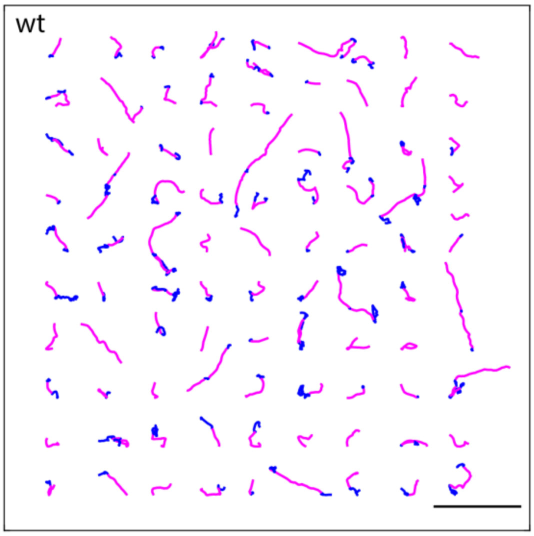


**Supplementary figure 4**: Compilation of HaloTag-MVI wild type (wt) tracks including runs (magenta) and pauses (blue) identified by recurrence analysis. Scale bar 5 µm.

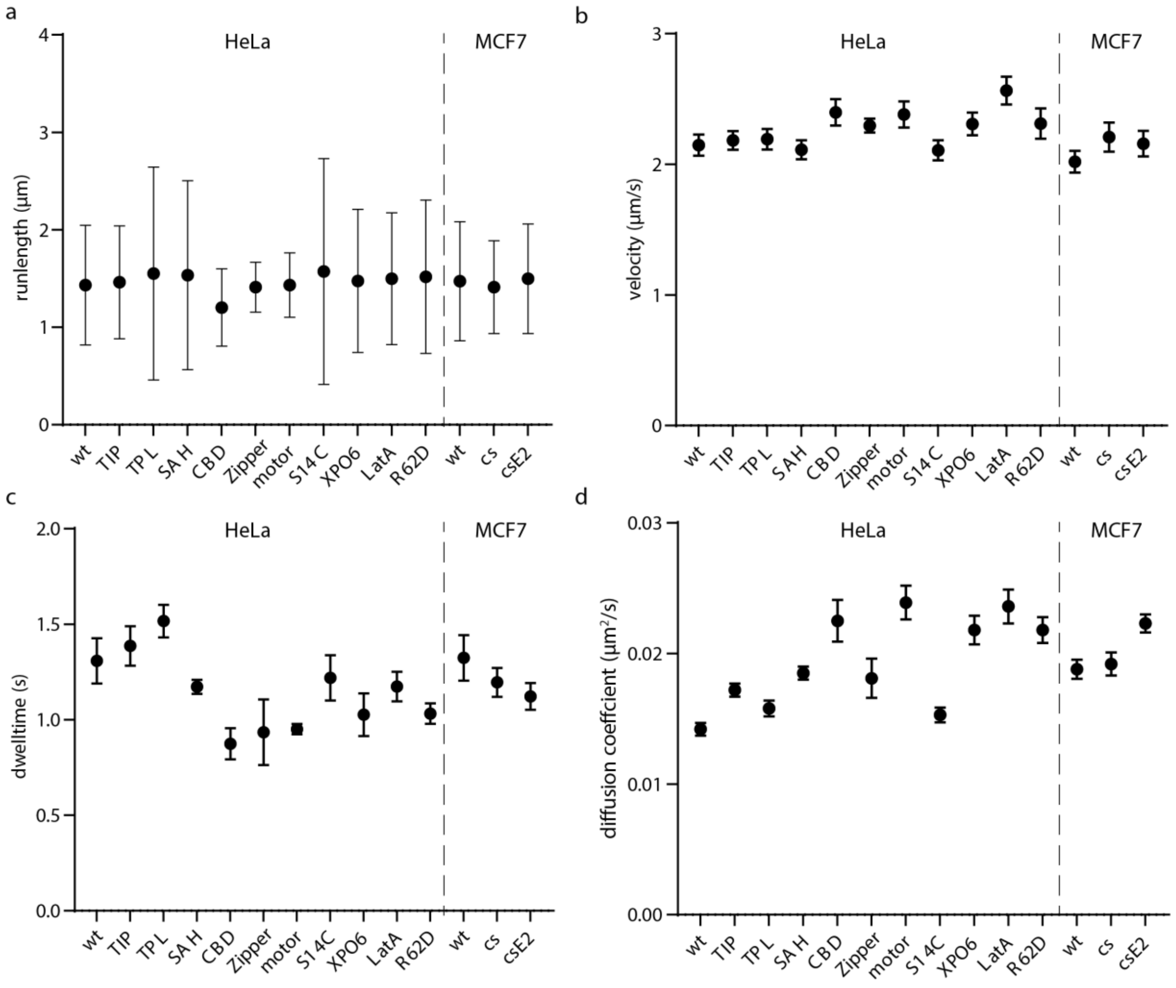


**Supplementary Figure 5:** Comparison of HaloTag-MVI track parameters at different conditions. Nomenclature and distributions is as in Supplementary Figures 4 to 17. (**a**) Median of runlength distributions. Error bars denote s.d. (**b**) Mean velocity determined by a Gaussian fit to velocity distributions. Error bars denote confidence interval of 95% (**c**) Average dwelltime determined by an exponential fit to dwelltime distributions. Error bars denote confidence interval of 95%. (**d**) Diffusion coefficient obtained from displacement distributions (Methods). Error bars denote sem.

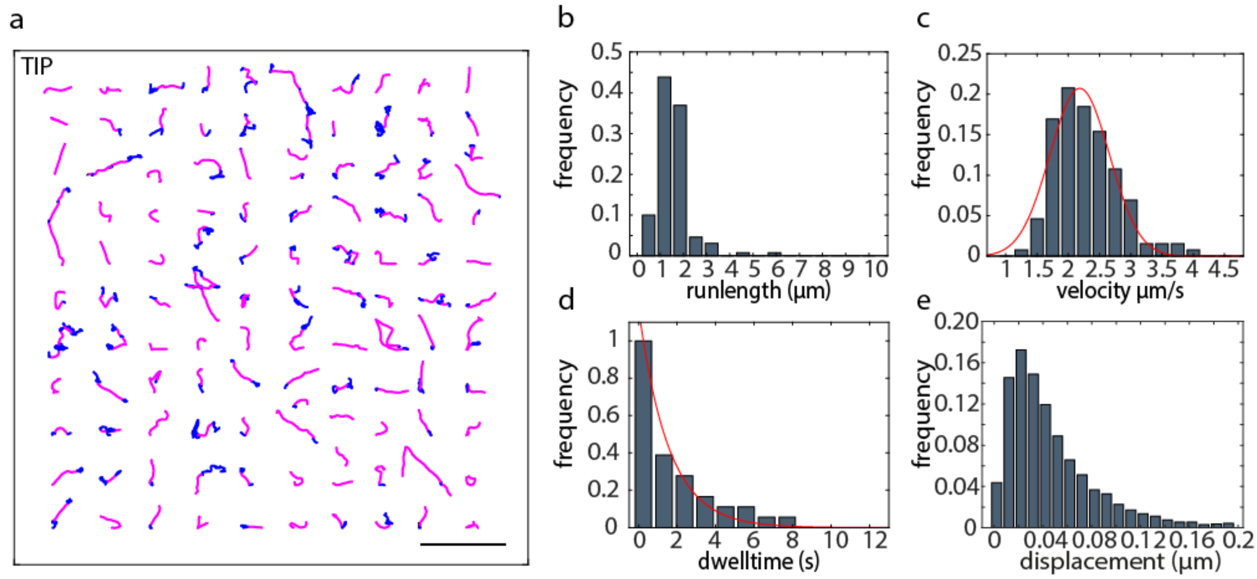


**Supplementary Figure 6:** Analysis of HaloTag-MVI motion in presence of 25 µM TIP. (**a**) HaloTag-MVI tracks including runs (magenta) and pauses (blue) identified by recurrence analysis. Scale bar 5 µm. Distributions of (**b**) runlength (1.5 ± 0.6 µm, median ± s.d.) and (**c**) velocity (2.2 ± 0.1 µm $s^{-1}$, mean ± s.d.) of MVI motion. (**d**) Distribution of dwell times (decay time: 1.4 ± 0.1 s, exponential fit ± confidence interval) and (**e**) step-size histogram (diffusion coefficient: 0.02 ± 0.001 $\mu m^2$ $s^{-1}$, diffusion coefficient ± sem, Methods) of MVI pauses.

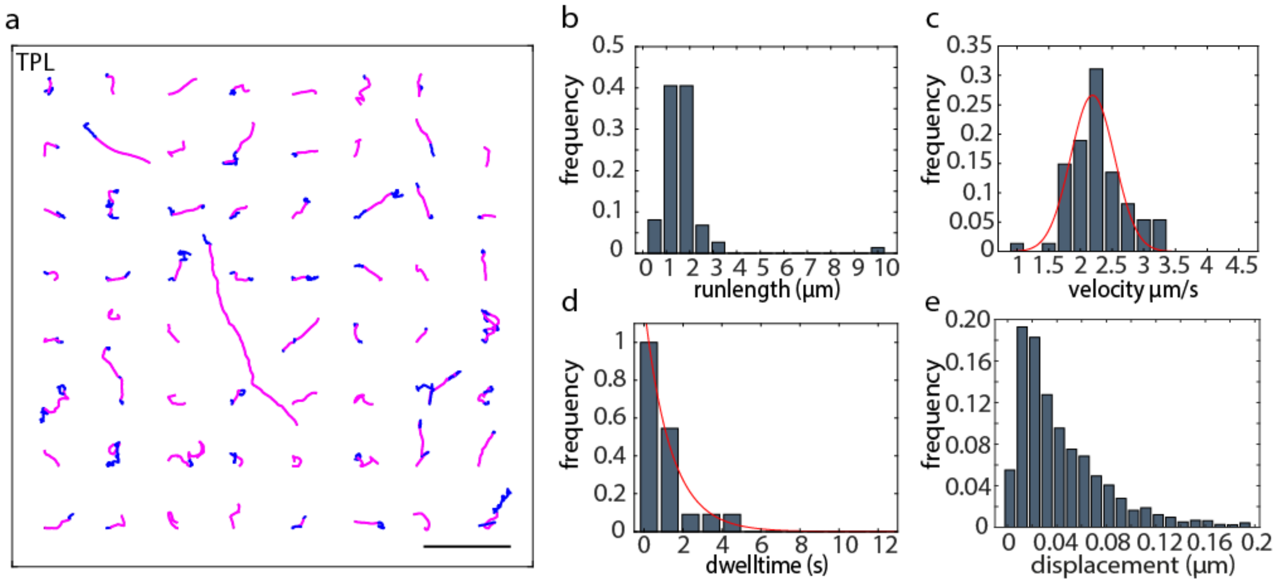


**Supplementary Figure 7:** Analysis of HaloTag-MVI motion in presence of 125 nM TPL. (**a**) HaloTag-MVI tracks including runs (magenta) and pauses (blue) identified by recurrence analysis. Scale bar 5 µm. Distributions of (**b**) runlength (1.6 ± 1.1 µm, median ± s.d.) and (**c**) velocity (2.2 ± 0.1 µm $s^{-1}$, mean ± s.d.) of MVI motion. (**d**) Distribution of dwell times (decay time: 1.5 ± 0.1 s, exponential fit ± confidence interval) and (**e**) step-size histogram (diffusion coefficient: 0.02 ± 0.001 $\mu m^2$ $s^{-1}$, diffusion coefficient ± sem, Methods) of MVI pauses.

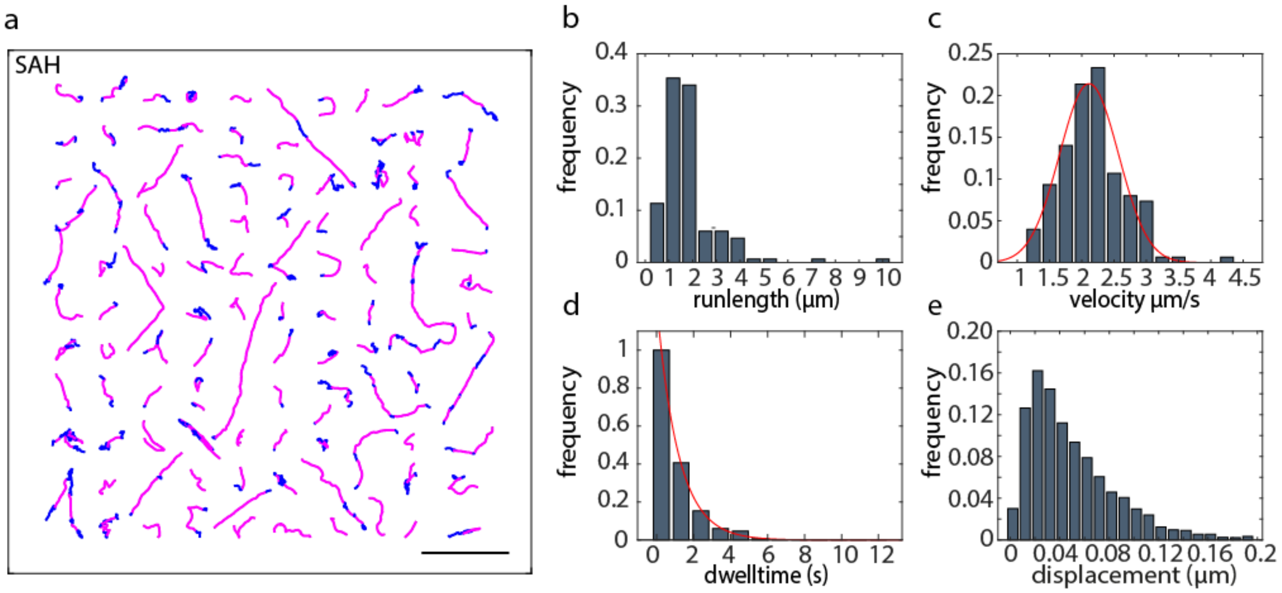


**Supplementary Figure 8:** Analysis of motion of HaloTag-MVI-SAH mutant. (**a**) HaloTag-MVI tracks including runs (magenta) and pauses (blue) identified by recurrence analysis. Scale bar 5 µm. Distributions of (**b**) runlength (1.5 ± 0.1 µm, median ± s.d.) and (**c**) velocity (2.1 ± 0.1 µm $s^{-1}$, mean ± s.d.) of MVI motion. (**d**) Distribution of dwell times (decay time: 1.2 ± 0.1 s, exponential fit ± confidence interval) and (**e**) step-size histogram (diffusion coefficient: 0.02 ± 0.001 µ$m^2$ $s^{-1}$, diffusion coefficient ± sem, Methods) of MVI pauses.

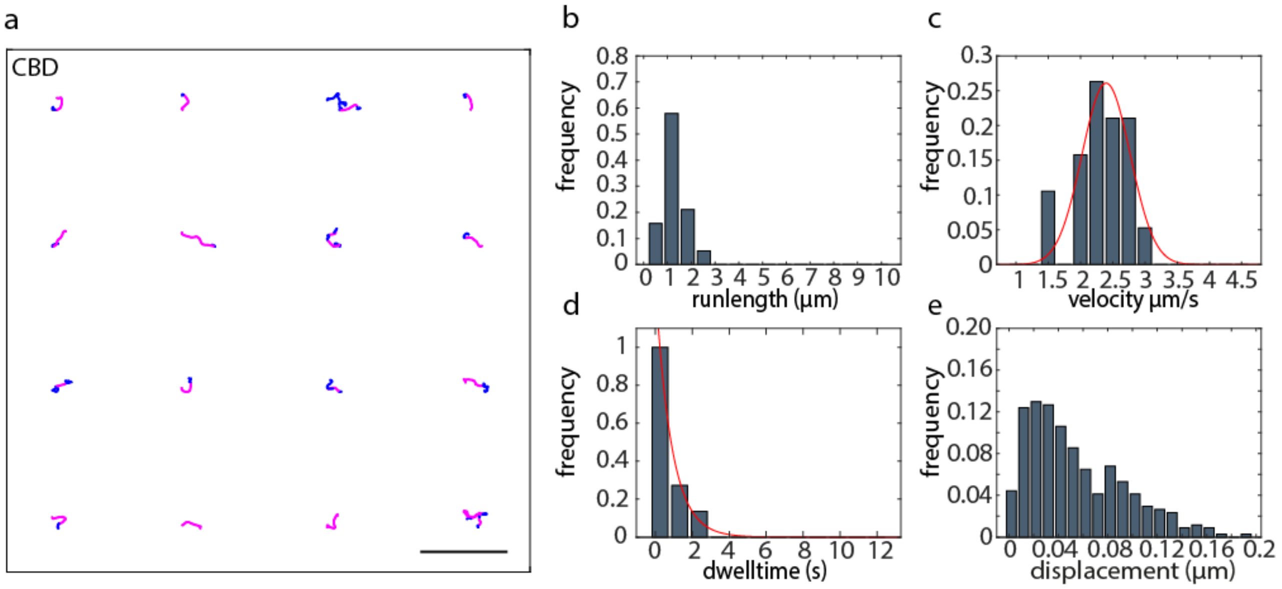


**Supplementary Figure 9:** Analysis of motion of HaloTag-MVI-CBD deletion constructs (**a**) HaloTag-MVI tracks including runs (magenta) and pauses (blue) identified by recurrence analysis. Scale bar 5 µm. Distributions of (**b**) runlength (1.2 ± 0.4 µm, median ± s.d.) and (**c**) velocity (2.4 ± 0.1 µm $s^{-1}$, mean ± s.d.) of MVI motion. (**d**) Distribution of dwell times (decay time: 0.9 ± 0.1 s, exponential fit ± confidence interval) and (**e**) step-size histogram (diffusion coefficient: 0.02 ± 0.002 µ$m^2$ $s^{-1}$, diffusion coefficient ± sem, Methods) of MVI pauses.

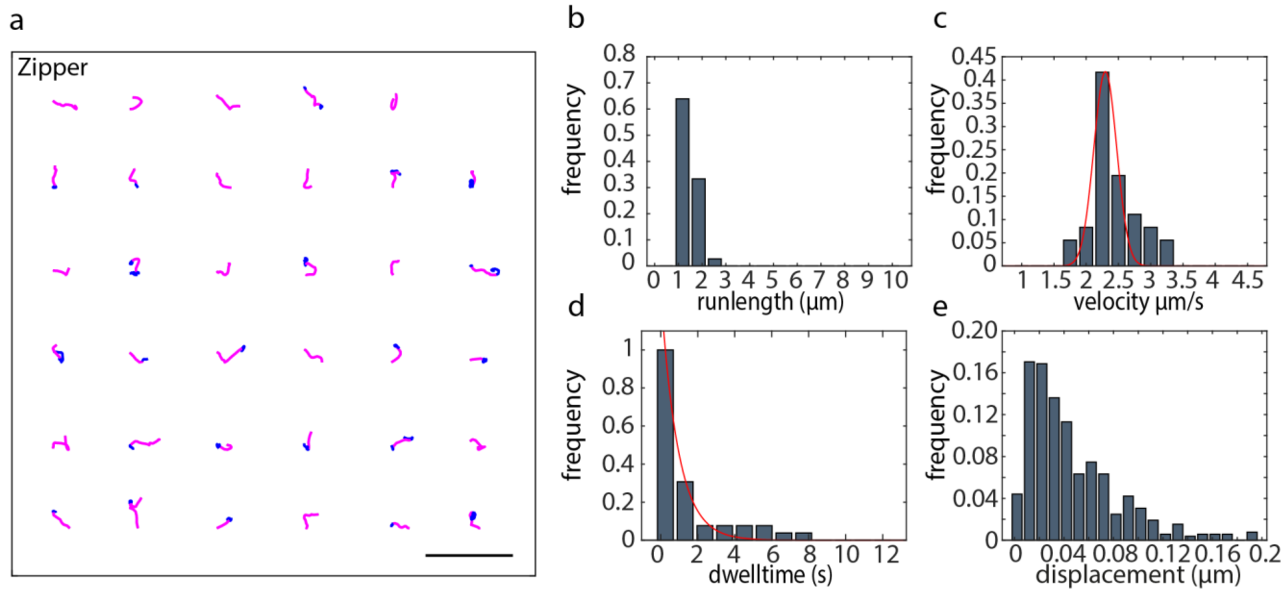


**Supplementary Figure 10:** Analysis of motion of HaloTag-MVI-Zipper deletion construct. (**a**) HaloTag-MVI tracks including runs (magenta) and pauses (blue) identified by recurrence analysis. Scale bar 5 µm. Distributions of (**b**) runlength (1.4 ± 0.3 µm, median ± s.d.) and (**c**) velocity (2.3 ± 0.1 µm $s^{-1}$, mean ± s.d.) of MVI motion. (**d**) Distribution of dwell times (decay time: 0.9 ± 0.2 s, exponential fit ± confidence interval) and (**e**) step-size histogram (diffusion coefficient: 0.02 ± 0.002 $\mu m^2$ $s^{-1}$, diffusion coefficient ± sem, Methods) of MVI pauses.

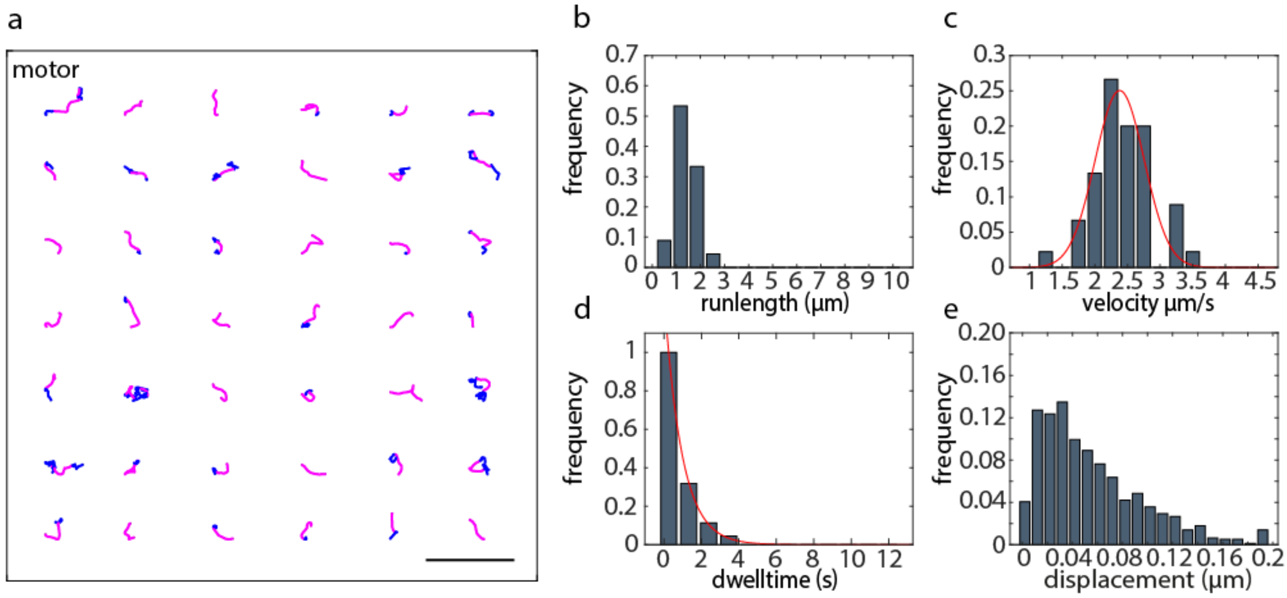


**Supplementary Figure 11:** Analysis of motion of HaloTag-MVI-motor deletion construct. (**a**) HaloTag-MVI tracks including runs (magenta) and pauses (blue) identified by recurrence analysis. Scale bar 5 µm. Distributions of (**b**) runlength (1.4 ± 0.3 µm, median ± s.d.) and (**c**) velocity (2.4 ± 0.1 µm $s^{-1}$, mean ± s.d.) of MVI motion. (**d**) Distribution of dwell times (decay time: 1.0 ± 0.1 s, exponential fit ± confidence interval) and (**e**) step-size histogram (diffusion coefficient: 0.02 ± 0.001 $\mu m^2$ $s^{-1}$, diffusion coefficient ± sem, Methods) of MVI pauses.

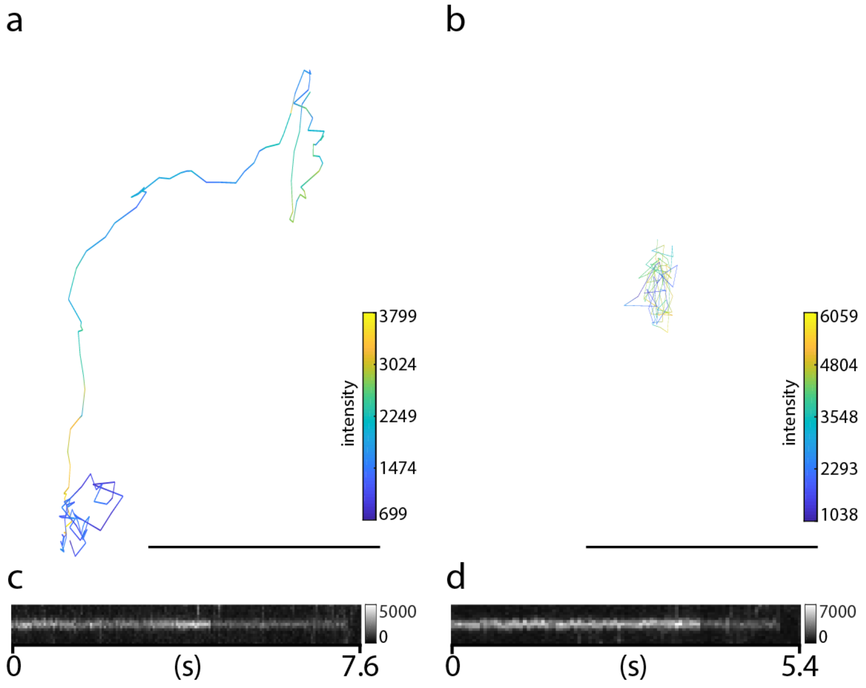


**Supplementary Figure 12:** Fluorescence intensity analysis of MVI tracks. (**a**) and (**b**) Tracks of MVI molecules. Color map indicates the intensity level. Scale bar 2 µm. (**c**) and (**d**) Kymographs with center position running along the tracks displayed in (a) and (b). Two consecutive bleaching steps can be observed.

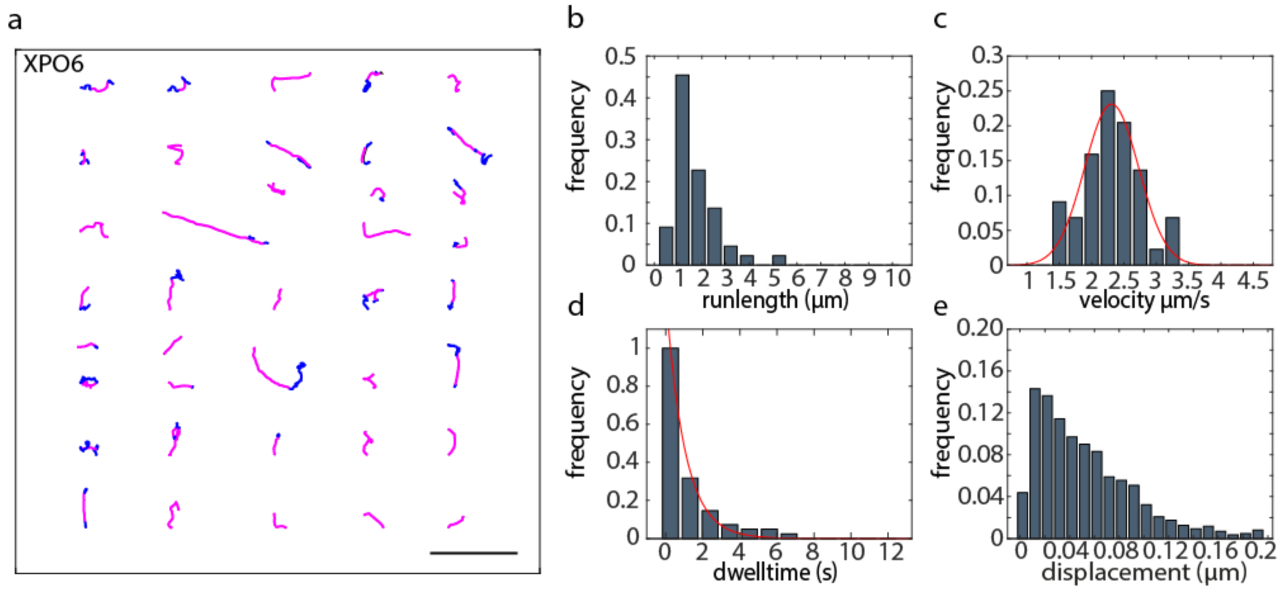


**Supplementary Figure 13:** Analysis of HaloTag-MVI motion in presence of conditions altering nuclear actin (XPO6). (**a**) HaloTag-MVI tracks including runs (magenta) and pauses (blue) identified by recurrence analysis. Scale bar 5 µm. Distributions of (**b**) runlength (1.5 ± 0.7 µm, median ± s.d.) and (**c**) velocity (2.3 ± 0.1 µm $s^{-1}$, mean ± s.d.) of MVI motion. (**d**) Distribution of dwell times (decay time: 1.0 ± 0.1 s, exponential fit ± confidence interval) and (**e**) step-size histogram (diffusion coefficient: 0.02 ± 0.001 µm$^2$ $s^{-1}$, diffusion coefficient ± sem, Methods) of MVI pauses.

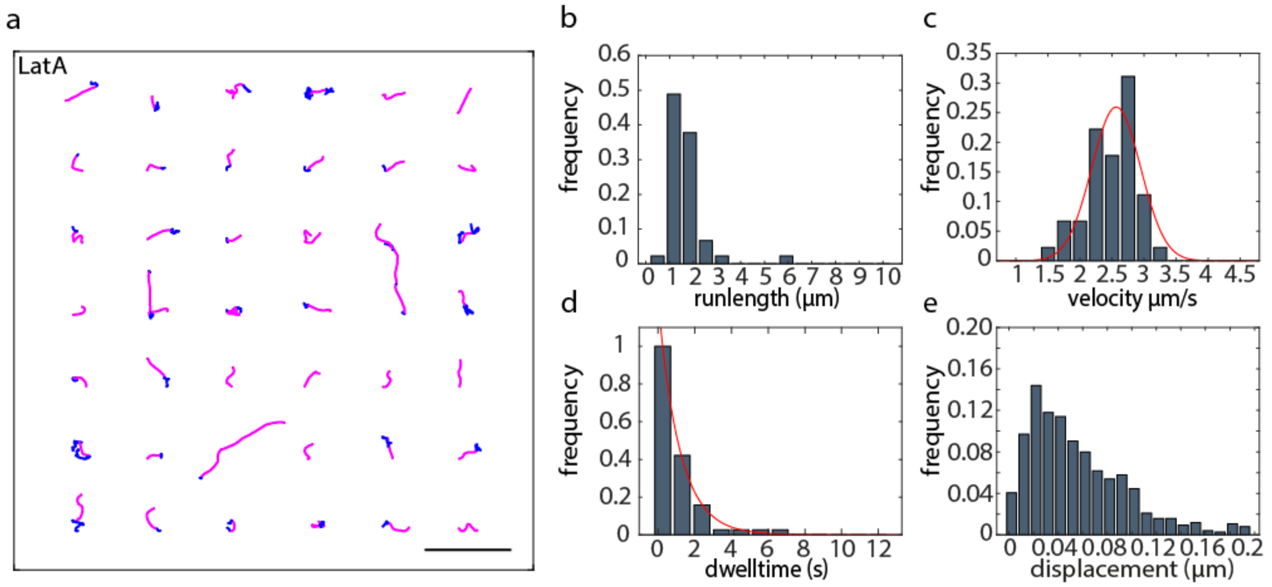


**Supplementary Figure 14:** Analysis of HaloTag-MVI motion in presence of conditions altering nuclear actin (LatA). (**a**) HaloTag-MVI tracks including runs (magenta) and pauses (blue) identified by recurrence analysis. Scale bar 5 µm. Distributions of (**b**) runlength (1.5 ± 0.7 µm, median ± s.d.) and (**c**) velocity (2.6 ± 0.1 µm $s^{-1}$, mean ± s.d.) of MVI motion. (**d**) Distribution of dwell times (decay time: 1.2 ± 0.1 s, exponential fit ± confidence interval) and (**e**) step-size histogram (diffusion coefficient: 0.02 ± 0.001 $\mu m^2$ $s^{-1}$, diffusion coefficient ± sem, Methods) of MVI pauses.

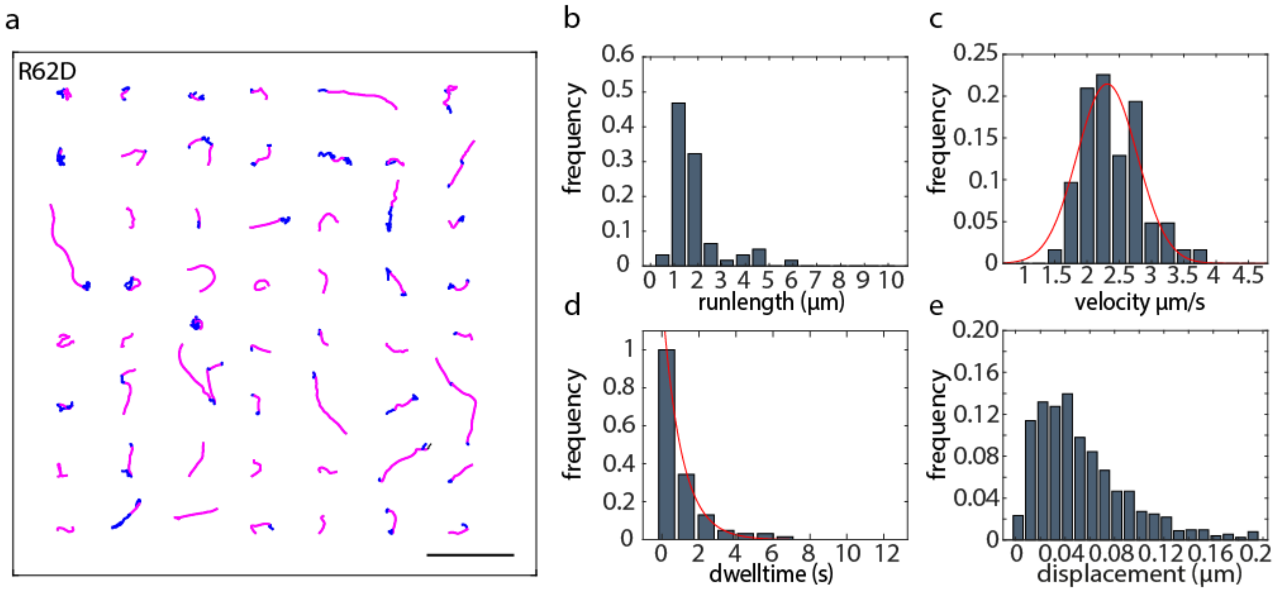


**Supplementary Figure 15:** Analysis of HaloTag-MVI motion in presence of conditions altering nuclear actin (R62D). (**a**) HaloTag-MVI tracks including runs (magenta) and pauses (blue) identified by recurrence analysis. Scale bar 5 µm. Distributions of (**b**) runlength (1.5 ± 0.8 µm, median ± s.d.) and (**c**) velocity (2.3 ± 0.1 µm $s^{-1}$, mean ± s.d.) of MVI motion. (**d**) Distribution of dwell times (decay time: 1.0 ± 0.1 s, exponential fit ± confidence interval) and (**e**) step-size histogram (diffusion coefficient: 0.02 ± 0.001 $\mu m^2$ $s^{-1}$, diffusion coefficient ± sem, Methods) of MVI pauses.

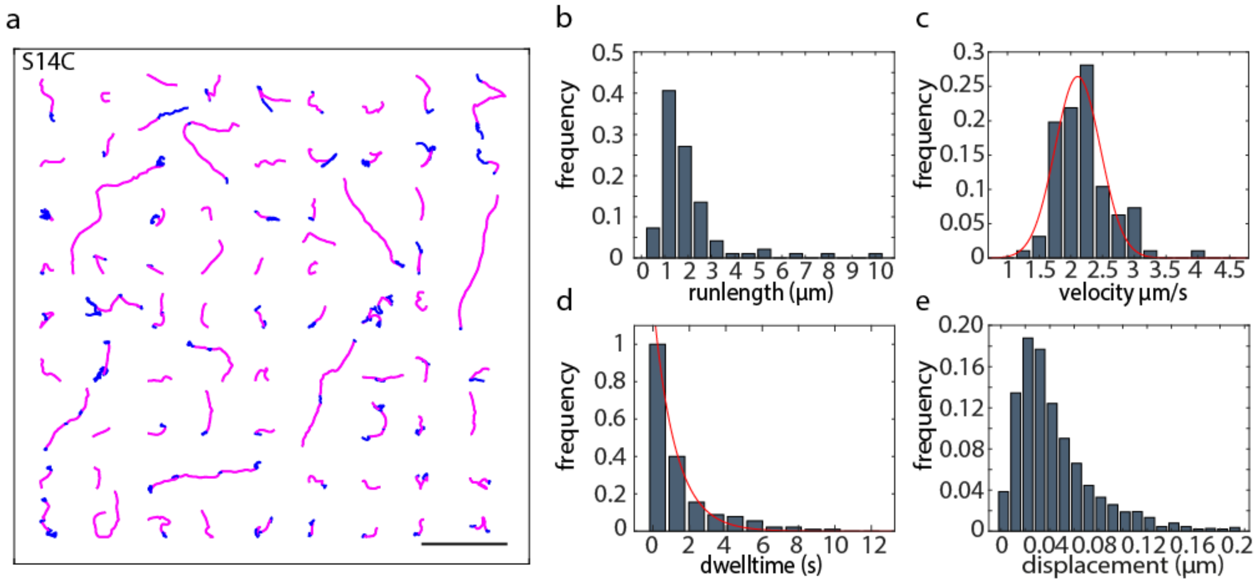


**Supplementary Figure 16:** Analysis of HaloTag-MVI motion in presence of conditions altering nuclear actin (S14C). (**a**) HaloTag-MVI tracks including runs (magenta) and pauses (blue) identified by recurrence analysis. Scale bar 5 µm. Distributions of (**b**) runlength (1.6 ± 1.2 µm, median ± s.d.) and (**c**) velocity (2.1 ± 0.1 µm $s^{-1}$, mean ± s.d.) of MVI motion. (**d**) Distribution of dwell times (decay time: 1.2 ± 0.1 s, exponential fit ± confidence interval) and (**e**) step-size histogram (diffusion coefficient: 0.02 ± 0.001 $\mu m^2$ $s^{-1}$, diffusion coefficient ± sem, Methods) of MVI pauses.

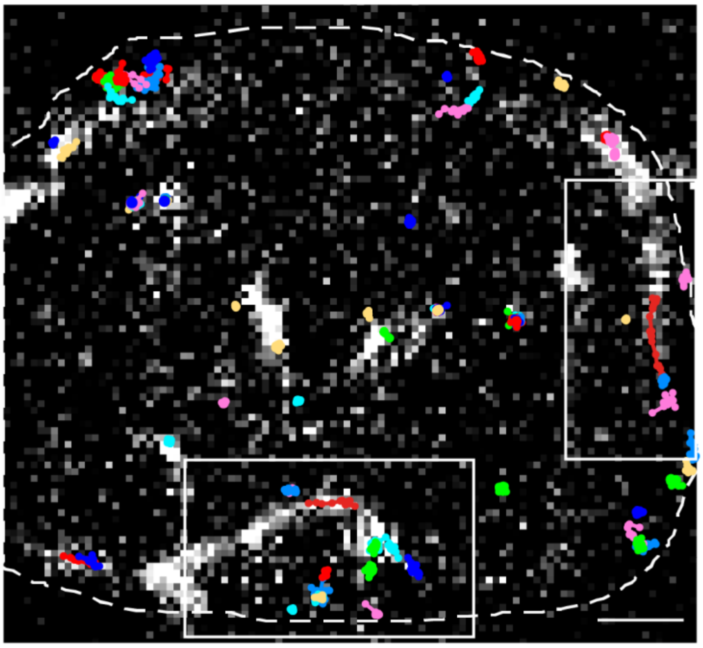

**Supplementary figure 17:** Co-localization of SiR-HaloTag-MVI tracks (colored lines) and nuclear actin chromobody (white) in HeLa cells. Boxes indicate tracks shown in Figure 3. Scale bar 2 µm.

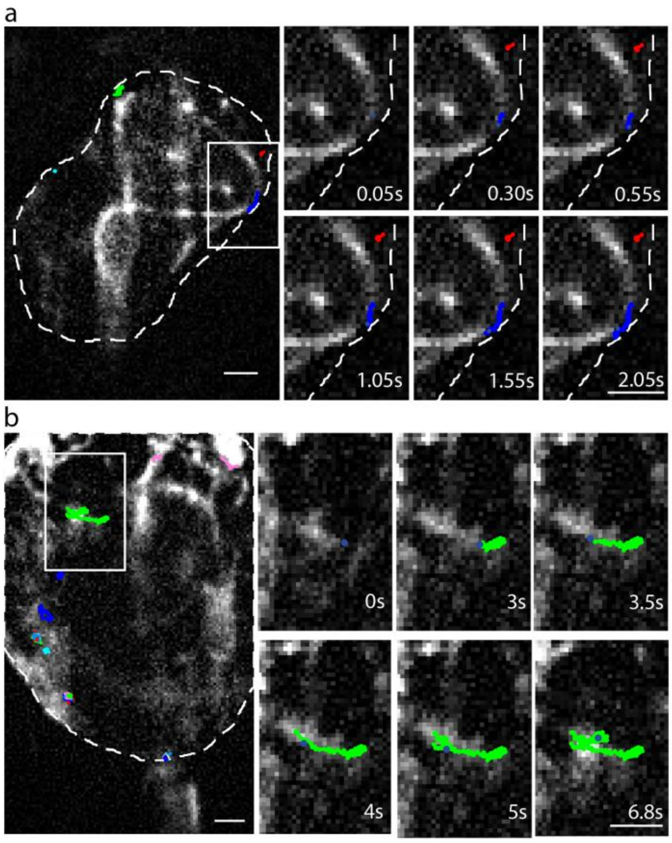


**Supplementary figure 18:** (**a**) Co-localization of SiR-HaloTag-MVI (colored lines) and nuclear actin chromobody (white) in HeLa cells and chronological series of zoom (white box). Scale bar 2 µm. (**b**) Co-localization of SiR-HaloTag-MVI (different colours) and nuclear actin chromobody (white) in MCF7 cells and chronological series of zoom (white box). Scale bar 2 µm.

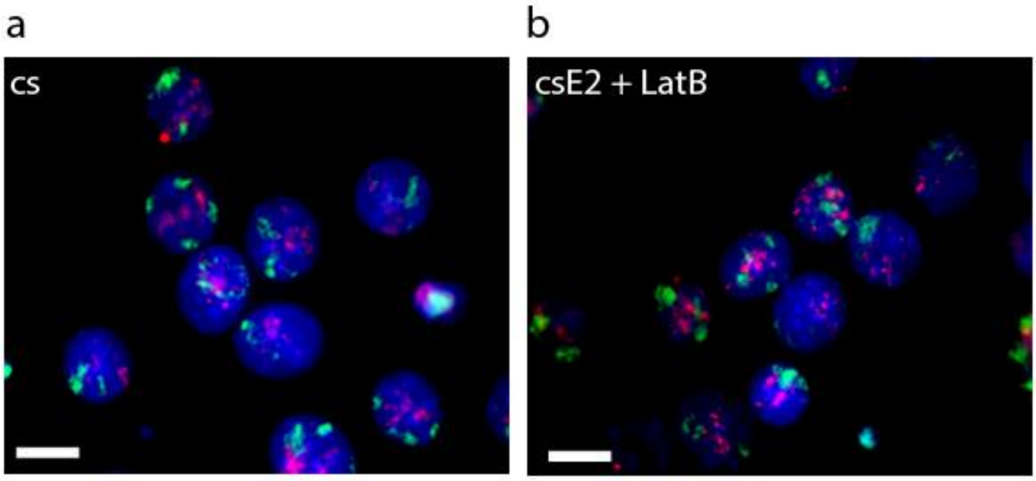


**Supplementary Figure 19:** Chromosome PAINT of chromosome 2 (green) and 21 (magenta) in MCF7 cells (**a**) after hormone depletion by growing cells in charcoaled stripped FCS (cs) or (**b**) after stimulation of transcription with β-estradiol and presence of acting filament destabilizing drug LatB (csE2 + LatB). Scale bar is 10 µm.

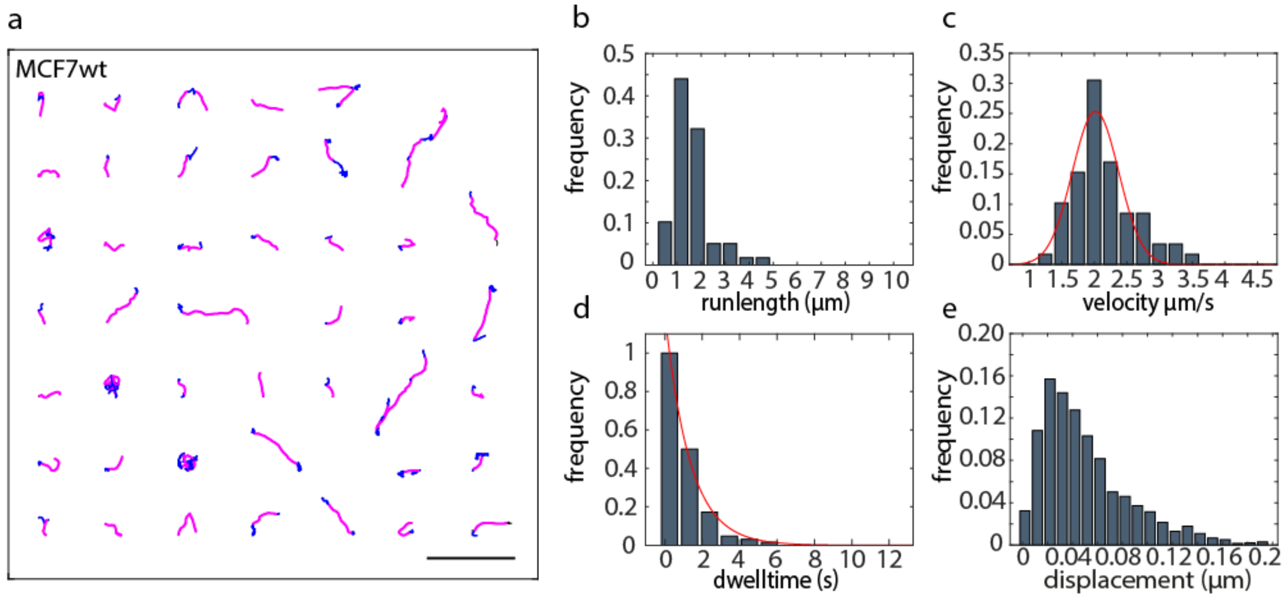


**Supplementary Figure 20:** Analysis of HaloTag-MVI motion in MCF7 (wt). (**a**) HaloTag-MVI tracks including runs (magenta) and pauses (blue) identified by recurrence analysis. Scale bar 5 µm. Distributions of (**b**) runlength (1.5 ± 0.6 µm, median ± s.d.) and (**c**) velocity (2.0 ± 0.08 µm $s^{-1}$, mean ± s.d.) of MVI motion. (**d**) Distribution of dwell times (decay time: 1.3 ± 0.1 s, exponential fit ± confidence interval) and (**e**) step-size histogram (diffusion coefficient: 0.02 ± 0.001 $µm^2$ $s^{-1}$, diffusion coefficient ± sem, Methods) of MVI pauses.

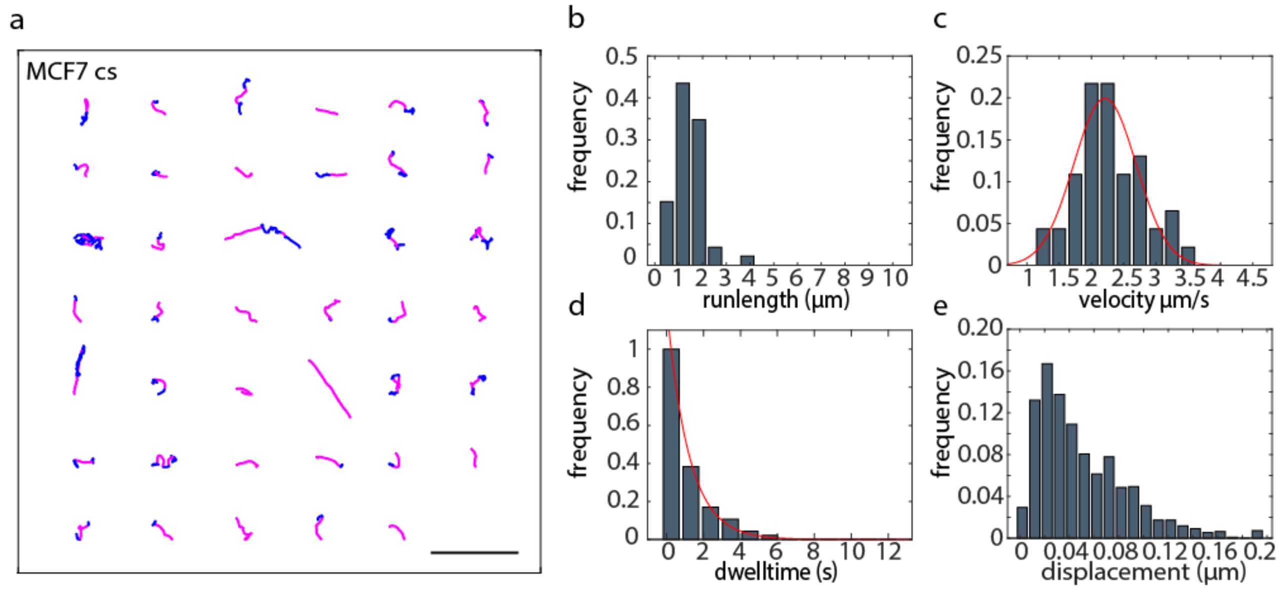


**Supplementary Figure 21:** Analysis of HaloTag-MVI motion in MCF7 cells treated 48h with charcoal stripped FCS (cs). (**a**) HaloTag-MVI tracks including runs (magenta) and pauses (blue) identified by recurrence analysis. Scale bar 5 µm. Distributions of (**b**) runlength (1.4 ± 0.5 µm, median ± s.d.) and (**c**) velocity (2.2 ± 0.1 µm $s^{-1}$, mean ± s.d.) of MVI motion. (**d**) Distribution of dwell times (decay time: 1.2 ± 0.1 s, exponential fit ± confidence interval) and (**e**) step-size histogram (diffusion coefficient: 0.02 ± 0.001 $µm^2$ $s^{-1}$, diffusion coefficient ± sem, Methods) of MVI pauses.

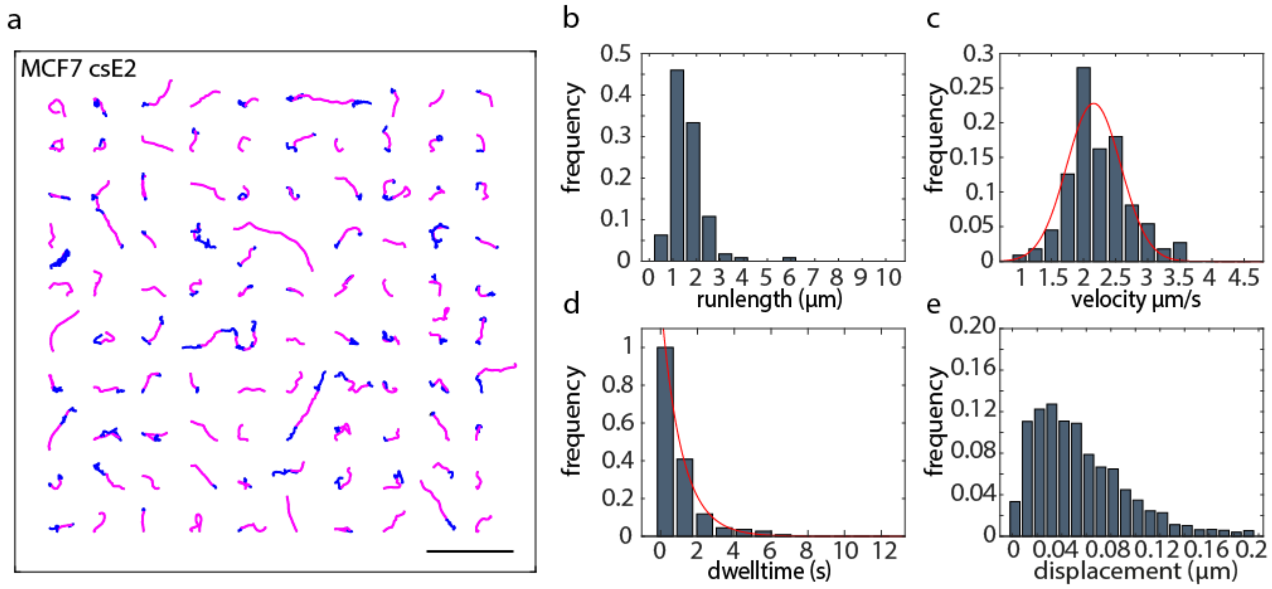


**Supplementary Figure 22:** Analysis of HaloTag-MVI motion in MCF7 cells treated 48h with charcoal stripped FCS (cs) and stimulation with β-estradiol (E2). (**a**) HaloTag-MVI tracks including runs (magenta) and pauses (blue) identified by recurrence analysis. Scale bar 5 µm. Distributions of (**b**) runlength (1.5 ± 0.6 µm, median ± s.d.) and (**c**) velocity (2.2 ± 0.1 µm $s^{-1}$, mean ± s.d.) of MVI motion. (**d**) Distribution of dwell times (decay time: 1.1 ± 0.1 s, exponential fit ± confidence interval) and (**e**) step-size histogram (diffusion coefficient: 0.02 ± 0.001 $\mu m^2 s^{-1}$, diffusion coefficient ± sem, Methods) of MVI pauses.

| Cluster No. | 1 | 2 | 3 | 4 |
|---|---|---|---|---|
| Dissociation rate | 0.01 | 0.14 | 0.7 | 7 |
| interval (1/s) | 0.14- | 0.7- | 7- | 70- |
| state sepctrum weight | (0.25 ± 0.06) % | (1.1 ± 0.1) % | (10 ± 0.6) % | (88 ± 0.7) % |
| event spectrum weight | (21 ± 2.6) % | (19 ± 2.1) % | (24 ± 0.9) % | (35 ± 1.6) % |

**Supplementary Table 1:** Positions and amplitudes of MVI dissociation rate clusters. Manually assigned dissociation rate intervals corresponding to a dissociation rate cluster are shown in row 2. Spectral weight is given by the sum over all amplitudes in one cluster. Errors denote s.d. obtained from 500 resampling runs.

| | | #cells | total #events | #motion | motion #events | normalized frequency | normalized sqrt(#motion) |
|---|---|---|---|---|---|---|---|
| HeLa | wt | 81 | 484857 | 92 | 1762 | 1.00 | 0.10 |
| | TIP | 104 | 795202 | 112 | 1769 | 0.61 | 0.06 |
| | TPL | 55 | 382691 | 63 | 1100 | 0.79 | 0.10 |
| | SAH | 56 | 471180 | 104 | 2510 | 1.47 | 0.14 |
| | CBD | 39 | 268357 | 16 | 195 | 0.20 | 0.05 |
| | zip | 45 | 325672 | 35 | 424 | 0.36 | 0.06 |
| | motor | 40 | 376680 | 42 | 522 | 0.38 | 0.06 |
| | S14C | 55 | 536111 | 76 | 1716 | 0.88 | 0.10 |
| | XPO6 | 67 | 365389 | 39 | 666 | 0.50 | 0.08 |
| | LatA | 57 | 285972 | 41 | 577 | 0.56 | 0.09 |
| | R62D | 84 | 558446 | 54 | 904 | 0.45 | 0.06 |
| MCF7 | wt | 44 | 275739 | 45 | 871 | 1.00 | 0.15 |
| | cs | 76 | 284164 | 40 | 566 | 0.63 | 0.10 |
| | csE2 | 77 | 452567 | 100 | 1584 | 1.11 | 0.11 |

**Supplementary Table 2:** Statistics of MVI track frequency calculation. To account for varying HaloTag-MVI expression levels in stable cell populations the total number of detection events (total #events) was used as normalization factor. Individual MVI tracks classified as directed motion (#motion) comprise a number of detection events (motion #events). The frequency of MVI motion was calculated by dividing motion #events by total #events and normalized to the corresponding wild type (wt). The error was estimated with the statistical counting error, sqrt(N), of the number of tracks classified as directed motion (#motion).

| Oligo sequences | Description |
|---|---|
| AAAAAA TCTAGA CCAAAAAAGAAGAGAAAGG | XbaI FL MVI FOR for insertion of MVIwt into LVTO |
| AAAAAA GGCGCGCC CTATTATTTCAACAGGTTCTG | AscI FL MVI REV for insertion of MVIwt into LVTO |
| AAAATCTAGACCAAAAAAGAAGAGAAAGGTAATGGAGG | XbaI_NLS_MVI_Motor_FOR for construction of LVTO-Halo_NLS_MVI_Motor |
| CTAGGGCGCGCCCTAAGCTCGATATTTTATTTTGTTTTTC | AscI_MVI_motor_REV for construction of LVTO-Halo_NLS_MVI_Motor |
| AAAATCTAGACCAAAAAAGAAGAGAAAGGTAGCAG | XbaI_NLS_MVI_CBD_FOR for construction of LVTO-Halo_NLS_MVI_CBD |
| AAAAGGCGCGCCCTATTTCAACAGGTTCTGCAGCAT | AscI_MVI_End_REV FOR for construction of LVTO-Halo_NLS_MVI_CBD |
| AAAATCTAGACCAAAAAAGAAGAGAAAGGTAATGGAAGATGGCAAGCCCGTG | XbaI_NLS_MVIzip_FOR for construction of LVTO-Halo_NLS_MVI_zip |
| AAAAGGCGCGCCTCATCTTTCGCC | AscI_MVIzip_REV for construction of LVTO-Halo_NLS_MVI_zip |
| AAAAGGATCCCCAAAAAAGAAGAGAAAGGTAGAGGATGGAAAACCCGTTTG | BamHI_NLS_MVI-SAH_FOR for construction of LVTO-Halo_NLS_MVI_SAH |
| AAAAGGCGCGCCCTACTTGAGCAGGTTCTGCAG | AscI_MVI_SAH_REV for construction of LVTO-Halo_NLS_MVI_SAH |
| TTCTTAGAATTCTCTAGAGTCGGATCCCTCGAGGGCGCGCCAATGTA | csRS_XbaI_BamHI_XhoI_AscI_fw for exchange of restriction cassete |
| TACATTGGCGCGCCCTCGAGGGATCCGACTCTAGAGAATTCTAAGAA | csRS_XbaI_BamHI_XhoI_AscI_rv for exchange of restriction casset |
| ATCGTCTAGAATGGCAGAAATCGGTACTGG | XbaI_Halo_fw for reinsertion of Halo-Tag |
| GCATGGATCCGCCGGAAATCTCGAGCGT | BamHI_Halo_rv for reinsertion of Halo-Tag |
| AATGGTCTCGAGATGACAGAATTTTTTCACGATTGT | XPO6_XhoI_fw for insertion of XPO6 into pIRES2_eGFP |
| ACCATTGAATTCCCCTAGAGCTTCACAGTGCCA | XPO6_EcoRI_rv for insertion of XPO6 into pIRES2_eGFP |

**Supplementary Table 3:** Oligo sequences (left) used for cloning of the different constructs (right).
